# SMARCAL1 is a candidate therapeutic target for ALT-positive tumors

**DOI:** 10.64898/2026.02.15.704070

**Authors:** Angelo Taglialatela, Jina Lee, Benura Azeroglu, Xiao Chen, Maria Rosaria Dello Stritto, Antoine Gouge, Tomas Lama-Diaz, Alina Vaitsiankova, Giuseppe Leuzzi, Filemon Dela Cruz, Zahra F Khan, Andrew L Kung, Petr Cejka, Eros Lazzerini Denchi, Jaewon Min, Alberto Ciccia

## Abstract

A significant subset of tumors, including over 50% of osteosarcomas—an aggressive bone malignancy affecting children, adolescents, and young adults—relies on alternative lengthening of telomeres (ALT), a telomerase-independent, DNA repair-based mechanism for telomere elongation. The overall 5-year survival rate for osteosarcoma patients is ∼65%, underlying the need to develop novel targeted therapies. Through the Cancer Dependency Map, we identify SMARCAL1, a DNA translocase previously shown to remodel stalled replication forks, as a top selective dependency factor in telomerase-negative tumors. Using a panel of ALT-positive and ALT-negative cancer cell lines, as well as osteosarcoma patient-derived xenograft cells, we confirm that ALT-positive cells are uniquely sensitive to the loss of SMARCAL1, whose depletion exacerbates ALT-dependent phenotypes and telomeric DNA damage. Notably, we demonstrate that suppressing ALT abrogates their dependency on SMARCAL1. Mechanistically, we show that SMARCAL1 loss leads to telomeric ssDNA accumulation in ALT-positive cells, dependent in part on DNA repriming mediated by the DNA primase/polymerase PRIMPOL. Moreover, SMARCAL1’s ssDNA annealing activity counteracts DNA unwinding by the BLM helicase, limiting telomeric ssDNA accumulation and DNA damage in ALT-positive cells. Importantly, SMARCAL1 depletion induces senescence in ALT-positive cancer cells, rendering them susceptible to treatment with senolytic agents. Together, these findings establish SMARCAL1 as a key regulator of ALT metabolism and highlight SMARCAL1 as a promising therapeutic target for ALT-positive tumors.

## Introduction

Tumor cells activate Telomere Maintenance Mechanisms (TMM) to preserve the stability and length of their telomeres during malignant proliferation. In most human tumors, the expression of the telomerase catalytic subunit *hTERT* is reactivated to counteract telomere shortening by adding new telomeric TTAGGG repeats to chromosomal ends at each round of genome duplication. A significant subset of human tumors employs an alternative TMM to elongate telomeres, known as the Alternative Lengthening of Telomeres (ALT) pathway (MacKenzie et al. 2021). Mutations in the ATRX/DAXX histone chaperone complex, which plays a key role in depositing the non-replicative histone variant H3.3 at pericentric and telomeric heterochromatin, are associated with the ALT status of cancer cells (Clatterbuck Soper and Meltzer 2023). ALT is triggered by spontaneous telomeric DNA damage, which promotes Break-Induced Replication (BIR), a homology-directed repair mechanism that facilitates the synthesis of large DNA segments to ensure telomere elongation (Loe et al. 2023; Sohn et al. 2023; O’Sullivan and Greenberg 2025). Numerous chromatin remodelers, and DNA replication and DNA Damage Response (DDR) factors are involved in ALT regulation, ensuring a coordinated balance of ALT activities that support the replicative immortality of cancer cells (Zhang and Zou 2020; Bhargava et al. 2022; Azeroglu et al. 2025). The ALT pathway depends on the DNA helicase BLM, which has been shown to unwind newly synthesized Okazaki fragments during telomere replication in ALT-positive cancer cells (Jiang et al. 2024; Lee et al. 2024), generating DNA intermediates that facilitate ALT through mechanisms that remain poorly understood. Deciphering the mechanisms that regulate ALT is therefore essential for developing effective treatments targeting this pathway in cancer.

CRISPR-Cas9 knockout (CRISPR-KO) screens have been employed to map genetic dependencies across more than 1,000 cancer cell lines (DepMap; https://depmap.org/portal/). Analyses of CRISPR-KO screens in pediatric cancer cell lines have suggested a potential essential function for the SNF2-family DNA translocase SMARCAL1 in a subset of pediatric tumors including osteosarcoma, a tumor type of mesenchymal origin frequently associated with an ALT-positive status and poor prognosis (Dharia et al. 2021; Harris and Hawkins 2022). Additional studies have shown that osteosarcoma cell lines and other ALT-positive cancer cells depend on the DNA translocase FANCM for their survival (Lu et al. 2019; Pan et al. 2019; Silva et al. 2019). FANCM has been reported to facilitate the repair of inter-strand crosslinks, the restart of stalled forks through fork reversal and the displacement of R-loop intermediates in response to replication stress (Gari et al. 2008; Xue et al. 2008; Deans and West 2009; Singh et al. 2009; Blackford et al. 2012; Wang et al. 2018). FANCM loss exacerbates telomere damage and promotes a hyper-ALT phenotype that is selectively toxic in cancer cells that already display features of ALT (Pan et al. 2017; Lu et al. 2019; Pan et al. 2019; Silva et al. 2019), with FANCM’s DNA translocase activity being required to attenuate telomeric replication stress in ALT-positive cancer cells (Lu et al. 2019; Pan et al. 2019; Silva et al. 2019). As such, FANCM loss or inhibition has been proposed to cause persistent accumulation of R-loops and stalled forks at telomeres of ALT-positive cells (Silva et al. 2019; Lu and Pickett 2022). Unlike FANCM, SMARCAL1 interacts with the single-strand DNA (ssDNA)-binding complex RPA to remodel stalled forks, promote fork reversal (Bansbach et al. 2009; Ciccia et al. 2009; Postow et al. 2009; Yuan et al. 2009; Yusufzai et al. 2009; Betous et al. 2012; Ciccia et al. 2012), and facilitate the restart of DNA synthesis (Bansbach et al. 2009; Ciccia et al. 2009; Postow et al. 2009; Yusufzai et al. 2009; Betous et al. 2012; Ciccia et al. 2012; Betous et al. 2013; Taglialatela et al. 2017). SMARCAL1 also possesses ssDNA annealing activity on bubble DNA structures coated with RPA (Yusufzai and Kadonaga 2008; Halder et al. 2022), and displaces R-loop intermediates (Hodson et al. 2022). SMARCAL1 has been proposed to prevent replication fork instability at telomeres in cancer cells regardless of their TMMs (Poole et al. 2015; Poole and Cortez 2017). Further studies have suggested a more prominent or specialized function of SMARCAL1 at telomeres of ALT-positive cancer cells (Cox et al. 2016; Zhang et al. 2019a; Zhang et al. 2019b; Feng et al. 2020; Salgado et al. 2024; Dubois et al. 2025; Lee et al. 2025). Moreover, BLM-dependent enrichment of SMARCAL1 at telomeres of ALT-positive cancer cells (Zhang et al. 2023; Jiang et al. 2024) supports a potential specific functional role for SMARCAL1 in the context of ALT.

In this study, by conducting a pan-cancer analysis of the DepMap database, we reveal a robust inverse correlation between *hTERT* expression and SMARCAL1 dependency, uncovering *SMARCAL1* as a top selective dependency gene in telomerase-negative cancer cells. Through validation studies, we confirm the association between SMARCAL1 dependency and ALT-positive status in cancer cells. Specifically, we show that suppression of ALT activities following BLM loss or DAXX re-expression in *DAXX*-mutant ALT-positive cancer cells bypasses the requirement for SMARCAL1 in preserving telomere stability and maintaining cell fitness. Additionally, we demonstrate that SMARCAL1 plays a protective role at telomeres of ALT-positive cancer cells, at least in part, by suppressing the repriming of DNA synthesis and the consequent formation of ssDNA gaps mediated by the DNA primase/polymerase PRIMPOL (Bai et al. 2020; Quinet et al. 2020). Moreover, SMARCAL1’s ssDNA annealing activity limits BLM-dependent unwinding of RPA-coated DNA substrates *in vitro,* and this function may be crucial to mitigate the telomere stress promoted by BLM’s activity in ALT-positive cancer cells (Jiang et al. 2024). Loss of SMARCAL1 in ALT-positive cancer cells also leads to increased accumulation of senescent cells and enhances cellular vulnerability to senolytic agents. These findings highlight SMARCAL1 as a potential therapeutic target for ALT-positive cancers.

## Results

### Identification of ALT-associated genetic vulnerabilities

To identify ALT-associated genetic vulnerabilities, we initially classified >1,300 cell lines listed in the DepMap database as telomerase-positive or -negative, based on *hTERT* expression, as determined using available RNA-seq data (Supplementary Table 1). Although telomerase activity and ALT-positive status can coexist in the same tumor (Perrem et al. 2001; Umaru et al. 2023), our classification reveals a clear inverse correlation between telomerase expression and ALT positivity, confirming recent studies highlighting the predictive value of lack of *hTERT* expression as a biomarker of ALT (Wu et al. 2025). Specifically, we observed the highest frequency of telomerase-negative cases within cell lines derived from tumors of mesenchymal origin, including osteosarcoma and other tumor types previously associated with high ALT prevalence (MacKenzie et al. 2021) (Supplementary Table 1). Comparative analysis of CRISPR-KO screens across all DepMap cancer cell lines (DepMap 23Q4) uncovered significant genetic dependencies selectively associated with telomerase-negative status. From this analysis, *SMARCAL1* emerged as the top selective dependency gene in telomerase-negative cancer cell lines (Figure 1A). Consistently, *SMARCAL1* displayed a greater mean dependency score in tumor types with a higher frequency of telomerase-negative cases, including osteosarcoma and other ALT-associated tumors (Supplementary Figure 1, Supplementary Table 2).

**Figure 1.**
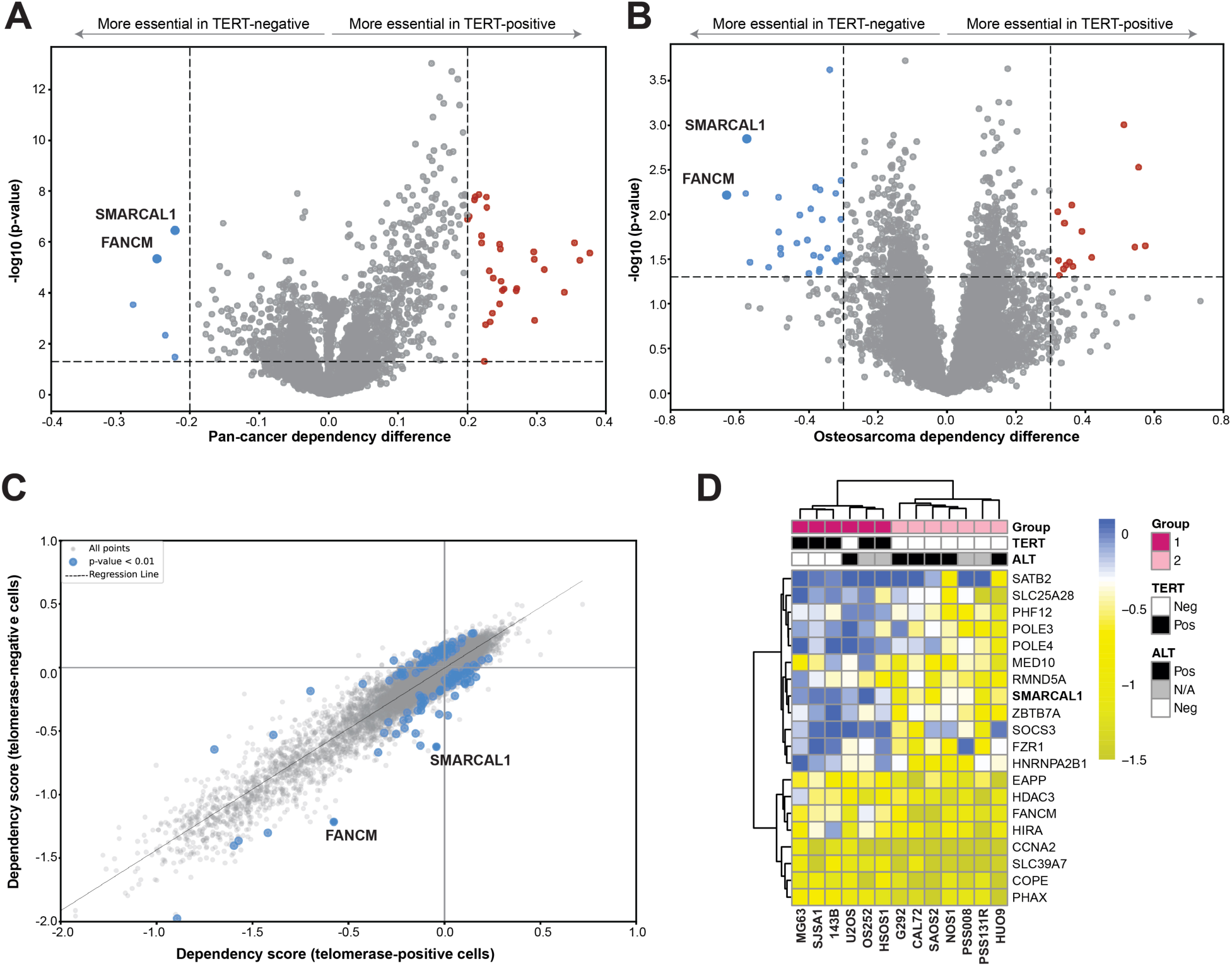
Analyses of cancer dependencies according to ALT status **(A-B)** Volcano plots showing differential gene dependencies between telomerase-negative and telomerase-positive cells, pan-cancer (1,009 cell lines) (A) and in osteosarcoma (13 cell lines) (B). Blue dots represent gene dependencies significantly enriched in telomerase-negative cells, red dots represent those enriched in telomerase-positive cells, and grey dots indicate genes without significant differences. The cut-offs for differential dependency scores and p-values were arbitrarily set. For clarity of presentation, two red dots were omitted from the graph. **(C)** Scatter plot of absolute gene effects in osteosarcoma, with telomerase-positive cells on the X-axis and telomerase-negative cells on the Y-axis. Each point represents a gene, with those along the diagonal having similar effects in both groups. Blue dots denote genes with significant differential dependency, while grey dots indicate no significant differences. Four grey dots were omitted from the graph for clarity of presentation. P-values were determined using an unpaired two-tailed Student’s t-test. **(D)** Clustered heatmap of the top 20 differential dependency genes (rows) in telomerase-positive and telomerase-negative osteosarcoma cells (columns), with *hTERT* and ALT status for each cell line indicated. The color scale represents dependency scores.

To mitigate potential confounding factors arising from the enrichment of cancer cell lines of mesenchymal origin observed after *hTERT* expression stratification, we conducted a differential analysis of gene dependencies within a single tumor type (osteosarcoma) stratified by *hTERT* expression. This analysis confirmed *SMARCAL1* among the top dependency genes in telomerase-negative osteosarcoma cell lines (Figure 1B, Supplementary Table 3). Moreover, in line with similar dependency analyses conducted using a smaller number of known ALT-positive cancer cell lines (Lu and Pickett 2022; Salgado et al. 2024; Dubois et al. 2025), we also observed *FANCM* as a significant dependency gene in telomerase-negative cancer cell lines (Figure 1A-B). Further corroborating the validity of our approach, we noted among the top significant differential dependencies additional genes previously implicated in the ALT pathway, including HIRA (Lee Lynskey et al. 2025), RMI1 (Loe et al. 2020) and TRIM28 (Kim et al. 2025; Tsai et al. 2025) (Supplementary Table 3). Comparison of the absolute gene dependency scores in telomerase-positive *vs* telomerase-negative osteosarcoma cell lines showed that SMARCAL1 loss displays more selective, although overall milder, detrimental effects towards telomerase-negative cells compared to the loss of FANCM (Figure 1C). Notably, clustering analysis of the top 20 most significant dependency genes successfully stratified osteosarcoma cell lines into two groups with largely distinct ALT/telomerase status (Figure 1D). Consistent with the above findings, *SMARCAL1* clustered together with genes whose loss did not cause significant cell fitness defects in telomerase-positive/ALT-negative cells (Figure 1D). This work uncovered *SMARCAL1* dependency as a specific biomarker of telomerase-negative/ALT-positive cancer cells.

### ALT-positive tumor models depend on SMARCAL1

To experimentally validate SMARCAL1 loss as a selective vulnerability in ALT-positive tumors, we performed transient knockdown of SMARCAL1 using RNA interference in a panel of osteosarcoma cell lines and assessed cell fitness relative to control siRNA-treated cells. A set of four distinct siRNAs targeting SMARCAL1 consistently impaired the fitness of ALT-positive cells, with no detrimental effects observed in ALT-negative cells (Figure 2A, Supplementary Figure 2A). Similarly, ALT-positive, but not ALT-negative, osteosarcoma patient-derived xenograft (PDX) cells exhibited fitness impairment upon SMARCAL1 depletion (Figure 2B, Supplementary Figure 2B).

**Figure 2.**
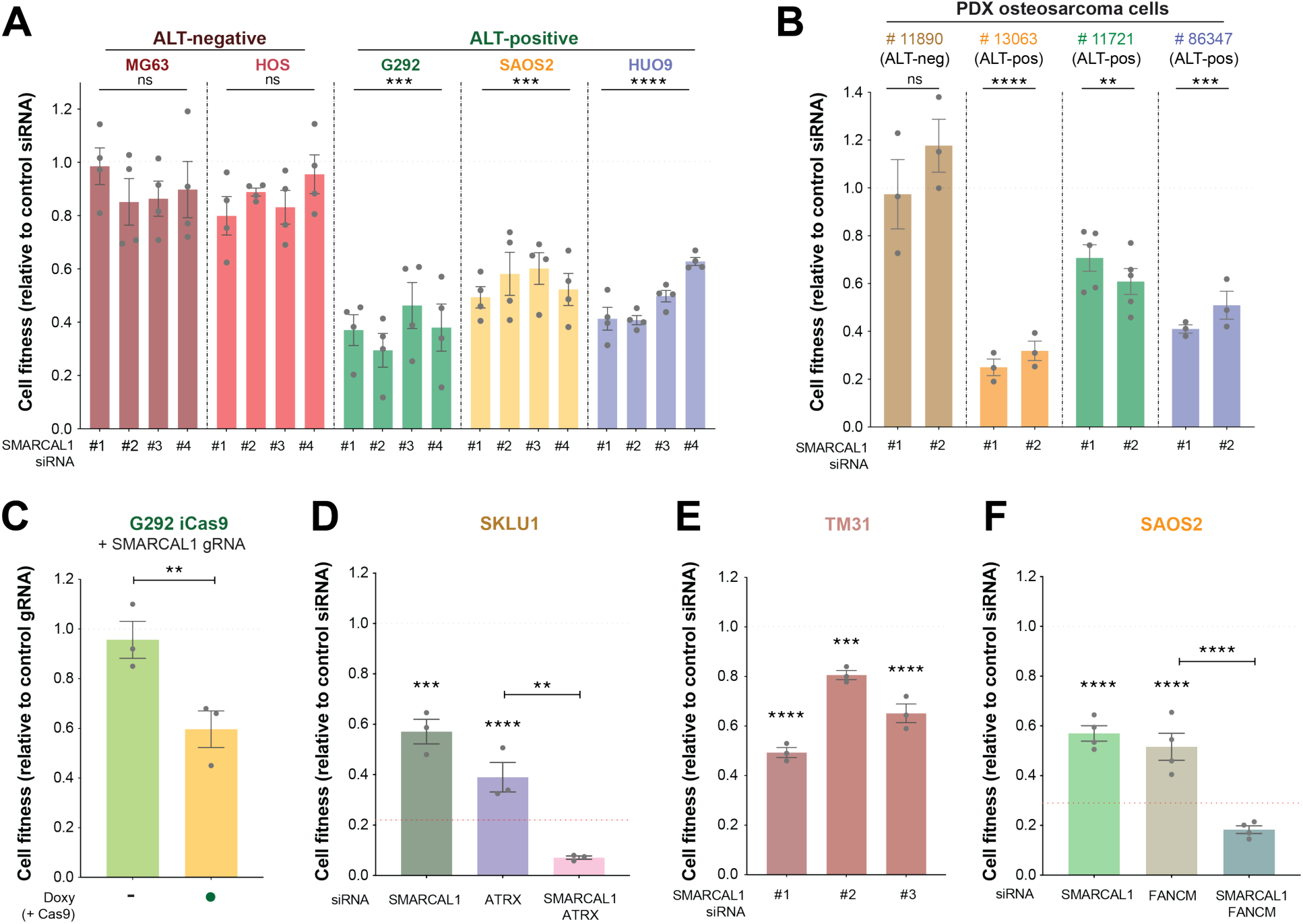
Dependency of ALT-positive osteosarcoma cells on SMARCAL1 **(A-F)** Cell fitness for the indicated cell lines transfected with the specified siRNAs, assessed using crystal violet staining to quantify viable cells. Data were normalized to cells treated with control (CTR) siRNA or gRNA following approximately 8 population doublings and represented as fold change relative to control (horizontal dashed grey line). The expected additive effect of dual gene targeting, calculated from single gene targeting data, is indicated by the dashed red line. Columns represent the mean ± SEM of independent biological replicates (dots). Statistical significance was determined using ordinary one-way ANOVA followed by Tukey’s multiple comparisons test. Significance levels are denoted as follows: ns (not significant), *p < 0.05, **p < 0.01, ***p < 0.001, ****p < 0.0001.

In line with the above findings, ALT-positive G292 cells expressing a doxycycline-inducible SpCas9 and a gRNA targeting *SMARCAL1* displayed reduced cell fitness upon *SMARCAL1* disruption (Figure 2C, Supplementary Figure 2D). Additionally, SMARCAL1 depletion impaired cell fitness in ALT-positive non-small cell lung cancer (SKLU1) and glioblastoma (TM31) cells (Figure 2D-E, Supplementary Figure 2C, E). Notably, SMARCAL1 depletion in ALT-positive cancer cells was associated with a marked increase of DNA damage response markers, including γH2AX and phospho-RPA2 (S4/S8) (Supplementary Figure 2D-E). To investigate the interplay between *SMARCAL1* and *FANCM* in the context of ALT, we analyzed the effects induced by their individual or combined loss in ALT-positive SAOS2 and OS384 osteosarcoma cells. Depletion or disruption of either SMARCAL1 or FANCM using siRNAs or CRISPR-Cas12a gRNAs resulted in similar levels of cell fitness impairment, while their combinatorial loss caused a synergistic reduction in cell fitness (Figure 2F, Supplementary Figure 2F-J), indicating non-epistatic functional roles for SMARCAL1 and FANCM in maintaining the viability of ALT-positive cancer cells. SMARCAL1 deficiency also reduced the fitness of SKLU1 cells (Figure 2D), an established ALT-positive cell line (Bryan et al. 1997; Brachner et al. 2006; Zhang et al. 2019a; Yadav et al. 2022) that retains functional ATRX and DAXX (Lovejoy et al. 2012). Interestingly, the combined depletion of SMARCAL1 and ATRX in SKLU1 cells further impaired cell fitness and enhanced DNA damage, as evidenced by the increase in phospho-RPA2 (S4/S8) levels (Figure 2D, Supplementary Figure 2E). These results suggest that *SMARCAL1* dependency is a general hallmark of ALT-positive cancer cells across diverse tumor types and genetic backgrounds.

### ALT suppression abrogates the dependency of ALT-positive cancer cells on SMARCAL1

To directly investigate the relationship between the ALT status and sensitivity to SMARCAL1 loss, we employed cellular systems amenable to ALT status modulation. Previous studies have shown that restoration of a fully functional ATRX/DAXX complex in ATRX/DAXX-deficient cells promptly suppresses ALT phenotypes in ATRX/DAXX-deficient models (Clynes et al. 2015; Napier et al. 2015; Mason-Osann et al. 2018; Yost et al. 2019). In our studies, we utilized a previously developed isogenic model of ALT suppression that relies on doxycycline-inducible DAXX expression in the G292 *DAXX*-mutant osteosarcoma cell line (G292iDAXX) (Yost et al. 2019). Using this cellular model, we first confirmed that SMARCAL1 depletion by siRNAs induced DNA damage, as determined by the accumulation of γH2AX and phospho-RPA2 (S4/S8) (Supplementary Figure 3A). Consistent with prior reports, wild-type DAXX expression in G292iDAXX cells following doxycycline treatment suppressed ALT-associated phenotypes, including the accumulation of DNA damage markers and the formation of ALT-associated PML bodies (APBs) (Supplementary Figure 3B-C). Similar to other ALT-positive cancer cells (Cox et al. 2016), G292iDAXX cells formed spontaneous SMARCAL1 foci that largely localized at telomeres, as determined by co-immunostaining for SMARCAL1 and the shelterin complex subunit TRF2 (Figure 3A). Additionally, G292iDAXX cells exhibited fitness impairment upon SMARCAL1 depletion, comparable to their parental counterpart (Figures 2A, C and 3B). Remarkably, suppression of ALT upon DAXX expression not only prevented the formation of SMARCAL1 foci (Figure 3A) but also suppressed both the cell fitness impairment and DNA damage accumulation caused by SMARCAL1 depletion (Figure 3B-C). Complementation experiments revealed that wild-type SMARCAL1 was able to rescue the DNA damage accumulation and the cell fitness impairment caused by SMARCAL1 depletion in G292 cells (Figure 3D-E, Supplementary Figure 3D-E). Two catalytically deficient mutants (R764Q and HD) (Ciccia et al. 2009; Ciccia et al. 2012), while clearly impaired relative to wild-type SMARCAL1, retained a reproducible ability to partially rescue the viability of SMARCAL1-deficient ALT-positive cells (Figure 3D-E, Supplementary Figure 3D-E). Notably, deletion of the amino-terminal RPA-binding region (DN_1-115_) (Bansbach et al. 2009; Taglialatela et al. 2017) similarly resulted in partial complementation, comparable to that observed with catalytic mutants (Figure 3D-E). However, combining the ΔN_1-115_ deletion with catalytic inactivation completely abolished rescue activity (Figure 3D-E). Finally, we tested whether loss of ZRANB3, a SMARCAL1-related DNA translocase of the SNF2 family, could recapitulate the phenotypes observed upon SMARCAL1 depletion in ALT-positive cells. However, siRNA-mediated depletion of ZRANB3 did not result in significant DNA damage accumulation and cell fitness defects in G292iDAXX cells (Supplementary Figure 3G-H), highlighting the non-redundant functions of SMARCAL1 in ALT-positive cells. Collectively, these results demonstrate that SMARCAL1 plays an essential, non-redundant role in cancer cells with a functional ALT pathway, dependent on both its DNA translocase activity and RPA-binding ability.

**Figure 3.**
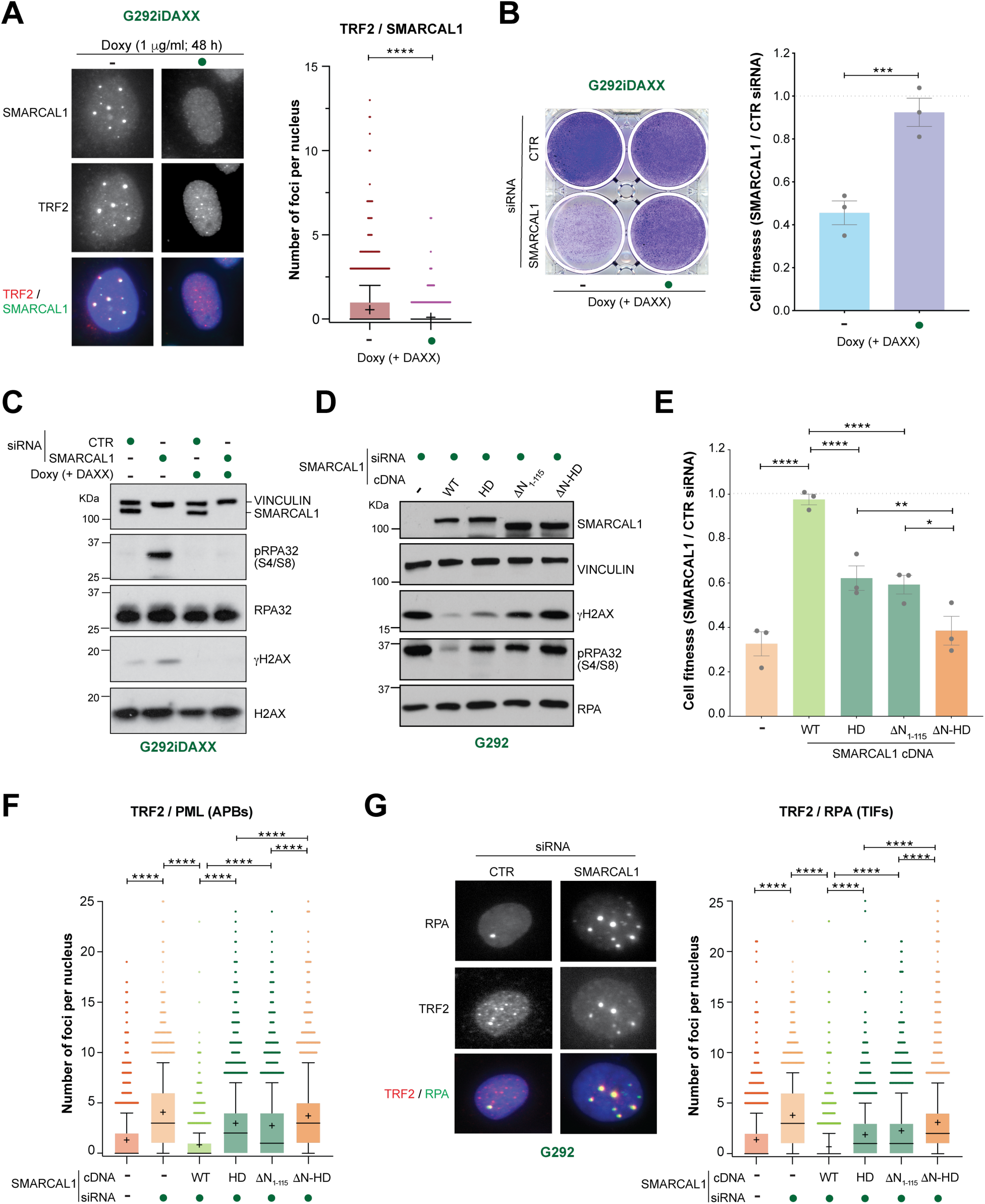
Cellular effects induced by SMARCAL1 loss in ALT-positive cancer cells (A) Representative images and quantification of SMARCAL1 foci per nucleus colocalizing with TRF2 in G292iDAXX cells, with or without doxycycline (doxy) treatment to express DAXX, as indicated. Data from 3 independent experiments are displayed as box-and-whisker plots, showing the median (line), mean (+), 1^st^ to 3^rd^ quartiles (colored box), whiskers extending to the 10^th^ and 90^th^ percentiles, and individual outlier points. Statistical significance was determined using ordinary one-way ANOVA followed by Tukey’s multiple comparisons test. (B) Representative images of G292iDAXX cells stained with crystal violet following transfection with the indicated siRNAs, with or without doxycycline treatment (Left panel). Quantification of the fitness of SMARCAL1 depleted cells (siRNA) relative to the control condition (CTR siRNA, horizontal dashed grey line) is shown (Right panel). Columns represent the mean ± SEM of independent biological replicates (dots). Statistical analysis and representation were conducted as in (A). **(C-D)** Detection by western blotting of the indicated proteins in whole cell extracts of G292iDAXX (C) and G292 (D) cells treated with the indicated siRNAs and/or expressing SMARCAL1 cDNA constructs, as shown. Vinculin is shown as loading control. **(E)** Quantification of the fitness of SMARCAL1-depleted (siRNA) G292 cells treated as in (D) relative to the control condition (CTR siRNA). Graphical representation and statistical analysis were conducted as in (B). **(F-G)** Quantification of the number of APBs (F), and representative images (G, left panel) and quantification of RPA32 (G, right panel) foci per nucleus colocalizing with TRF2 in G292 cells transfected with the indicated cDNA constructs and siRNAs. Data representation and statistical analysis were conducted as in (A). Significance levels are denoted as follows: ns (not significant), *p < 0.05, **p < 0.01, ***p < 0.001, ****p < 0.0001.

### SMARCAL1 confers telomere stability in ALT-positive cancer cells

Consistent with earlier findings (Cox et al. 2016), SMARCAL1 depletion in ALT-positive cancer cells led to telomeric DNA damage, as evidenced by an increase in the number of telomere dysfunction-induced foci (TIFs) marked by γH2AX foci colocalizing with TRF2 (Supplementary Figure 4A-B). Furthermore, SMARCAL1 depletion exacerbated the formation of APBs, as indicated by the enhanced colocalization of PML bodies with TRF2 (Figures 3F and 4A, Supplementary Figure 3F). Notably, ALT suppression in G292iDAXX cells upon DAXX expression fully rescued the telomeric DNA damage and the formation of APBs induced by SMARCAL1 depletion (Figure 4A-C). Interestingly, FANCM depletion in SAOS2 cells resulted in increased SMARCAL1 localization at telomeres (Supplementary Figure 4D) and further enhanced telomeric DNA damage when combined with SMARCAL1 loss (Supplementary Figure 4C). These findings suggest that hyper-activation of the ALT pathway resulting from FANCM loss (Lu et al. 2019; Silva et al. 2019) renders ALT-positive cells increasingly dependent on SMARCAL1 to mitigate telomere stress.

**Figure 4.**
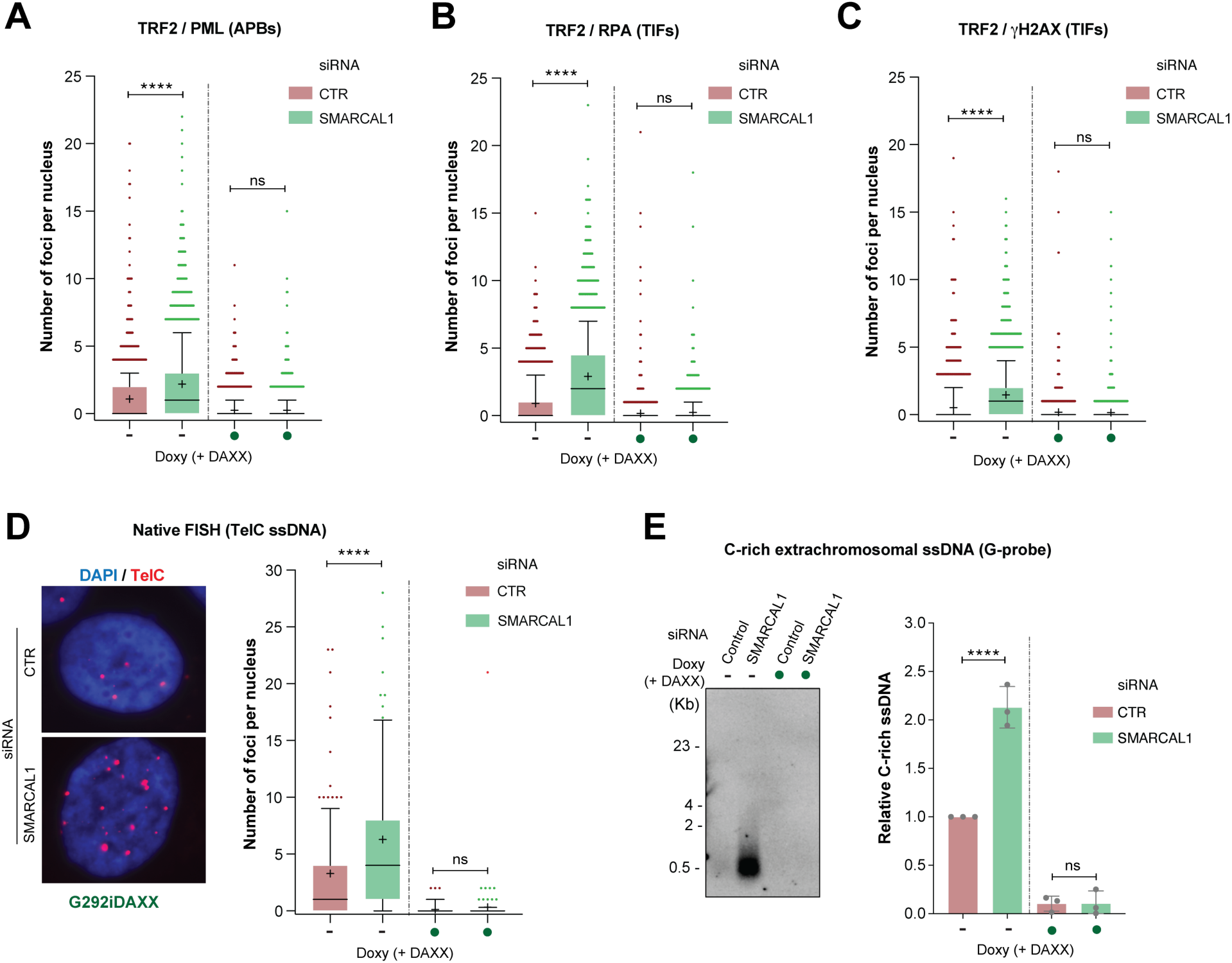
Analysis of telomeric DNA damage and ssDNA formation following SMARCAL1 loss in ALT-positive cancer cells **(A-C)** Quantification of the number of PML (A), RPA32 (B), and γH2AX (C) foci per nucleus colocalizing with TRF2 in G292iDAXX cells transfected with the indicated siRNAs, with or without doxycycline (doxy) treatment to express DAXX, as indicated. Data from 3 independent experiments are represented as box-and-whisker plots, showing the median (line), mean (+), 1^st^ to 3^rd^ quartiles (colored box), whiskers extending to the 10^th^ and 90^th^ percentiles, and individual outlier points. Statistical significance was determined using ordinary one-way ANOVA followed by Tukey’s multiple comparisons test. (D) Representative images of ssTelo and quantification of the number of telomeric ssDNA foci per nucleus in G292iDAXX cells transfected with the indicated siRNAs, with or without doxycycline treatment. Data representation and statistical analysis were conducted as in (A). (E) Detection of C-rich extrachromosomal telomeres by 4SET assay performed on the soluble fraction of G292iDAXX cells transfected with the indicated siRNAs with or without doxycycline treatment. Representative chemilumigram image (left panel) and quantification of C-rich telomeric DNA (right panel) using a (Digoxigenin)-labeled G-rich DNA probe are shown. Columns represent the mean ± SEM of independent biological replicates (dots). Statistical analysis was conducted as in (A). Significance levels are denoted as follows: ns (not significant), ****p < 0.0001.

To define the nature of the telomeric DNA damage induced by SMARCAL1 depletion in ALT-positive cells, we monitored the formation of RPA foci, a ssDNA marker, in G292 cells. SMARCAL1 depletion significantly increased RPA foci colocalizing with TRF2, an effect that was fully suppressed by DAXX expression (Figures 3G and 4B). Complementation with wild-type SMARCAL1 fully suppressed APB and TIF formation induced by depletion of endogenous SMARCAL1 (Figure 3F-G). Consistent with our ALT cell viability assays (Figure 3E), the DN_1-115_ and HD mutants each conferred partial suppression, an effect that was abolished in the double mutant, indicating that both RPA binding and DNA translocase activity are required for SMARCAL1 to restrain ALT activity (Figure 3F-G).

To directly monitor the formation of telomeric ssDNA in the above settings, we performed telomere FISH analysis in G292iDAXX cells under non-denaturing conditions (ssTelo) (Loe et al. 2020; Azeroglu et al. 2024). These studies revealed a significant increase in nuclear C-rich telomeric DNA foci upon SMARCAL1 depletion (Figure 4D). Additionally, SMARCAL1 loss triggered the accumulation of telomeric C-rich extrachromosomal ssDNA, as detected by 4SET analysis on the soluble fraction of G292iDAXX cells (Figure 4E), indicating that the ALT-dependent accumulation of telomeric ssDNA products is accentuated in the absence of SMARCAL1. Both nuclear and extrachromosomal telomeric ssDNA signals induced by SMARCAL1 depletion were suppressed by DAXX expression (Figure 4D-E). Together, these findings indicate that SMARCAL1 plays a critical role in maintaining telomere stability by limiting the accumulation of telomeric ssDNA, thus tightly regulating ALT metabolism in ALT-positive cancer cells.

### PRIMPOL-mediated DNA repriming promotes telomere instability in ALT-positive cancer cells upon SMARCAL1 loss

SMARCAL1-dependent fork reversal operates at stalled forks alternatively to the restart of DNA synthesis mediated by the DNA primase/polymerase PRIMPOL (Bai et al. 2020; Quinet et al. 2020). To determine whether the interplay between SMARCAL1 and PRIMPOL is relevant in the context of ALT-positive cells, we investigated whether modulation of PRIMPOL levels would influence the cellular effects induced by SMARCAL1 depletion. Notably, we observed that overexpression of wild-type PRIMPOL exacerbated the cell fitness defects induced by SMARCAL1 depletion in G292 cells (Figure 5A). Furthermore, PRIMPOL overexpression led to a further increase in RPA2 phosphorylation in SMARCAL1-depleted G292 cells (Supplementary Figure 5A). Unlike expression of wild-type PRIMPOL, expression of repriming (CH), catalytically dead (AxA), and RPA-binding (RBD) PRIMPOL mutants failed to aggravate the cell fitness defects induced by SMARCAL1 depletion in G292 cells (Figure 5B, Supplementary Figure 5B). These findings suggest that PRIMPOL’s DNA repriming activity can contribute to the accumulation of telomeric ssDNA in the context of ALT. However, depletion of endogenous PRIMPOL only partially suppressed RPA2 phosphorylation and telomeric ssDNA accumulation in SMARCAL1-depleted G292 cells (Figure 5C-D, Supplementary Figure 5C), suggesting that additional PRIMPOL-independent mechanisms contribute to telomeric ssDNA accumulation and DNA damage in the absence of SMARCAL1.

**Figure 5.**
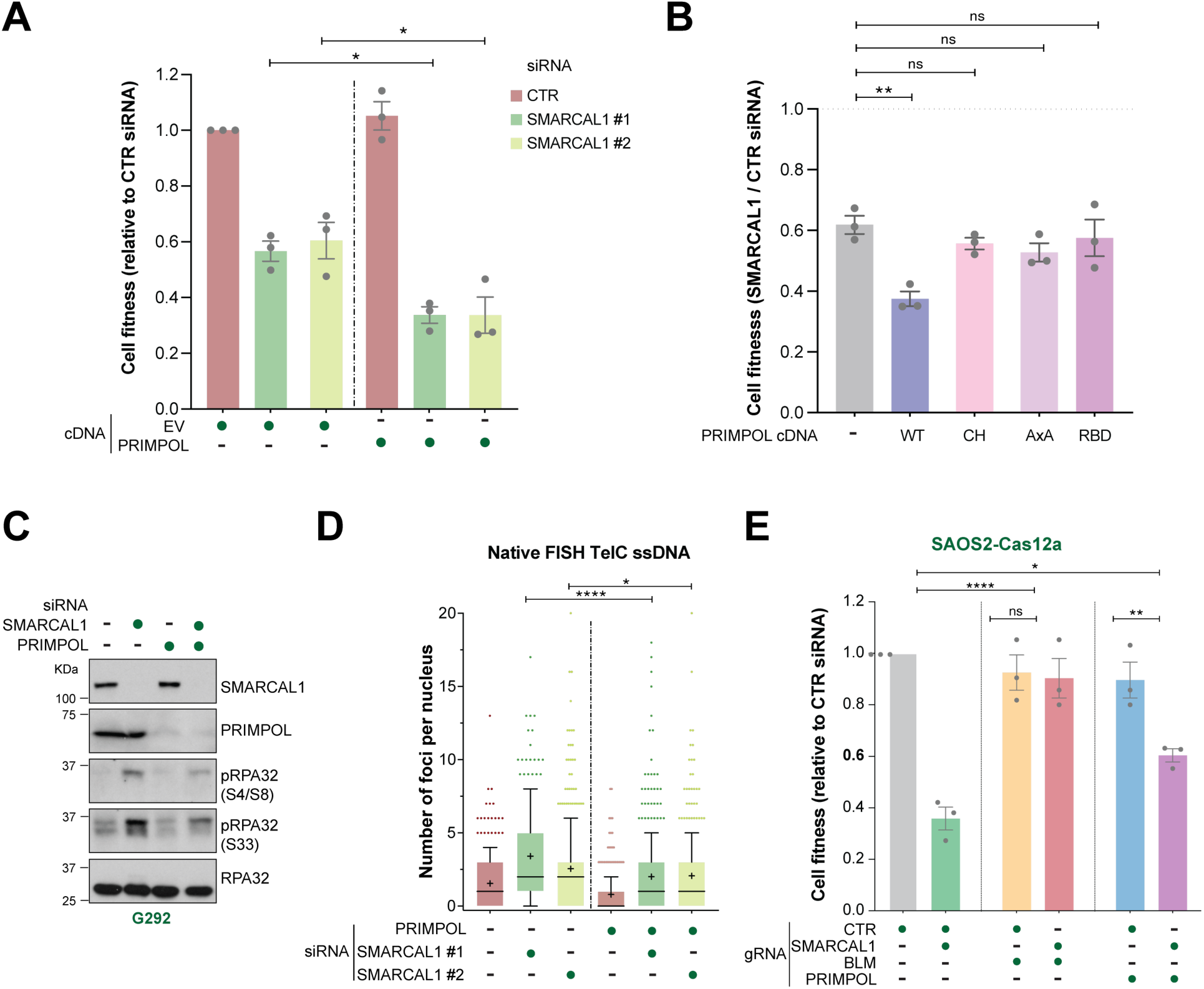
PRIMPOL-dependent cellular effects induced in ALT-positive cancer cells following SMARCAL1 loss **(A-B)** Quantification of the fitness of G292 cells transfected with the indicated cDNAs and siRNAs relative to the control condition (CTR siRNA or untreated). Columns represent the mean ± SEM of independent biological replicates (dots). Statistical significance was determined using ordinary one-way ANOVA followed by Tukey’s multiple comparisons test. **(C)** Detection by western blotting of the indicated proteins in whole cell extracts of G292 cells transfected with the indicated siRNAs. RPA32 is shown as a loading control. **(D)** Quantification of the number of ssTelo foci per nucleus in G292 cells transduced with the indicated siRNAs. Data are represented as box-and-whisker plots, showing the median (line), mean (+), 1^st^ to 3^rd^ quartiles (colored box), whiskers extending to the 10^th^ and 90^th^ percentiles, and individual outlier points. Data were obtained from 3 independent experiments. Statistical analysis was conducted as in (A-B). **(E)** Quantification of the fitness of SAOS2 cells expressing Cas12a and transduced with the indicated gRNAs, relative to the control conditions (CTR gRNA), as determined by cell counting. Columns represent the mean ± SEM of independent biological replicates (dots). Statistical analyses were conducted as in (D). Significance levels are denoted as follows: ns (not significant), *p < 0.05, **p < 0.01, ***p < 0.001, ****p < 0.0001.

### SMARCAL1 counteracts BLM activity at damaged telomeres

Multiple studies have demonstrated that the BLM helicase is critical for both the initiation and the maintenance of the ALT pathway (Stavropoulos et al. 2002; O’Sullivan et al. 2014; Sobinoff et al. 2017; Min et al. 2019; Zhang et al. 2019a; Loe et al. 2020; Zhang et al. 2021). In light of our findings, we sought to investigate the impact of BLM loss on the phenotypes observed in SMARCAL1-depleted ALT-positive cells and compare it to that of PRIMPOL loss. To this end, we employed Cas12a to disrupt either *BLM* or *PRIMPOL* together with *SMARCAL1* in ALT-positive cells (Supplementary Figure 6G). Interestingly, loss of BLM, but not PRIMPOL, significantly impaired SMARCAL1 enrichment at telomeres in ALT-positive cells (Figure 6D, Supplementary Figure 6A, D). Consistent with BLM’s established role in ALT, BLM depletion reduced APB formation in G292 cells and suppressed the elevated APB levels induced by SMARCAL1 loss (Supplementary Figure 6B). In addition, BLM loss fully suppressed the accumulation of telomeric damage induced by *SMARCAL1* disruption, as determined by monitoring RPA- and γH2AX-positive TIFs (Supplementary Figure 6C, E-F). In contrast, PRIMPOL loss induced a partial reduction of the accumulation of telomeric RPA foci induced by SMARCAL1 loss, without altering the formation of APBs (Supplementary Figure 6B-C). Consistently, PRIMPOL loss partially mitigated the cell fitness defects induced by SMARCAL1 loss in SAOS2 and G292 cells, while BLM loss fully suppressed the SMARCAL1-dependent growth defects in those cells (Figure 5E, Supplementary Figure 6G-H).

**Figure 6.**
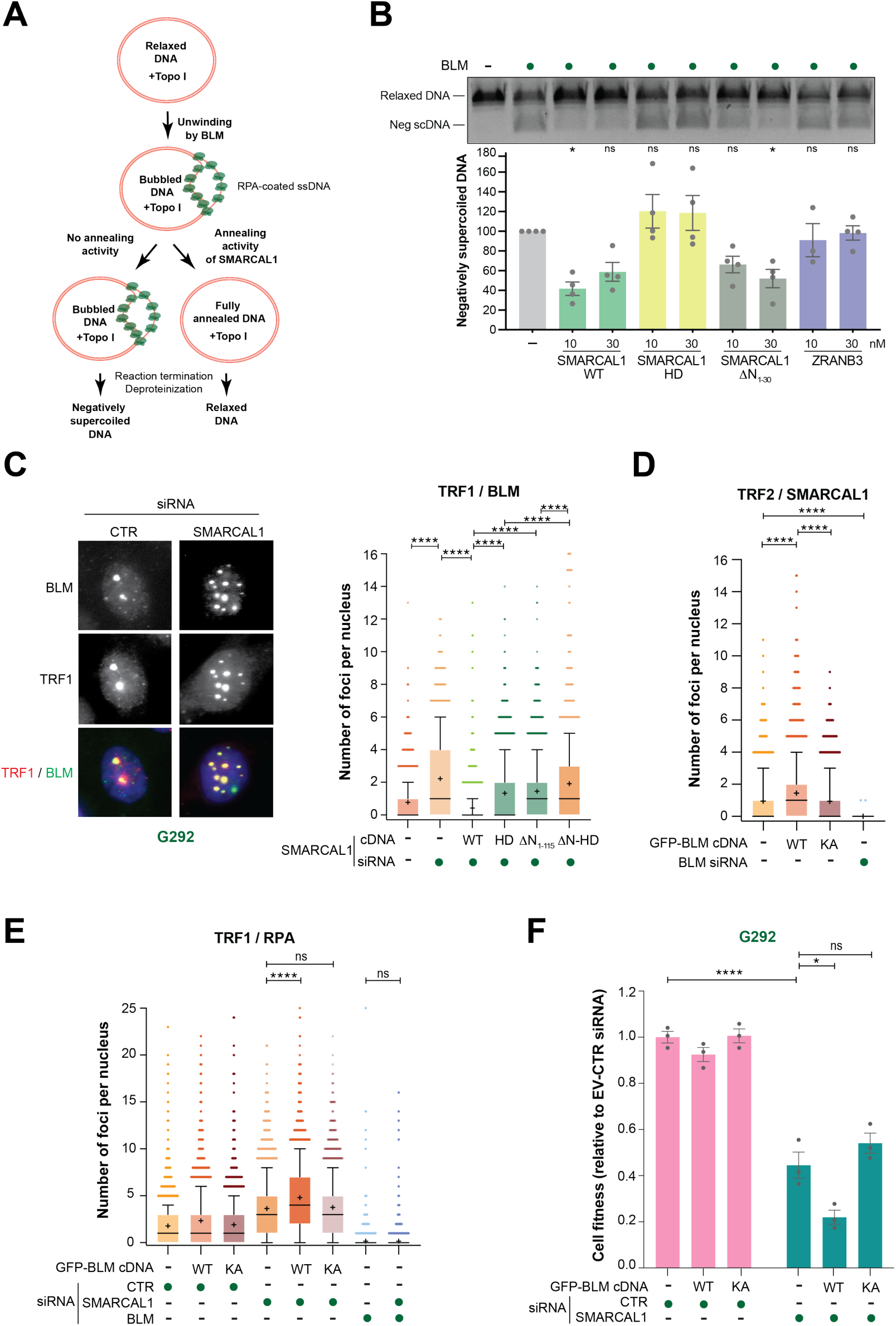
Antagonistic interplay between BLM and SMARCAL1 in ALT-positive cancer cells (A) Schematic of the plasmid-based, topoisomerase-coupled unwinding/annealing assay. (B) Activities of SMARCAL1, WT and mutants (HD, ΔN_1-30_), and ZRANB3 in topoisomerase-coupled annealing assays. Top, representative experiment. Bottom, quantification of supercoiled DNA levels relative to those generated by BLM alone. Columns represent the mean ± SEM of independent replicates (dots). Statistical significance was determined using ordinary one-way ANOVA followed by Tukey’s multiple comparisons test. (C) Representative images and quantification of BLM foci per nucleus colocalizing with TRF1 in G292 cells expressing the indicated cDNAs and transfected with the indicated siRNAs. Statistical analysis was conducted as in (B). **(D-E)** Quantification of SMARCAL1 foci per nucleus colocalizing with TRF2 (D), and RPA32 foci colocalizing with TRF1 (E) in G292 cells transfected with the indicated cDNAs and siRNAs. Data representation and statistical analysis were conducted as in (B). **(F)** Quantification of the fitness of G292 cells by cell counting upon transfection with the indicated siRNAs and cDNAs, relative to cells treated with an empty vector (EV) and a control siRNA (CTR siRNA) following approximately 8 population doublings. Columns represent the mean ± SEM of independent biological replicates (dots). Statistical analysis was conducted as in (B). Significance levels are denoted as follows: ns (not significant), *p < 0.05, **p < 0.01, ***p < 0.001, ****p < 0.0001.

To gain mechanistic insights into the functional interaction between BLM and SMARCAL1, we assessed their enzymatic activities. Using a topoisomerase-coupled plasmid-based assay (Yusufzai and Kadonaga 2008), we observed that BLM induced the formation of ssDNA bubbles stabilized by RPA, resulting in negatively supercoiled DNA (Figure 6A-B). Remarkably, addition of SMARCAL1 to the same reaction suppressed the formation of negatively supercoiled DNA by BLM, revealing SMARCAL1’s ability to counteract BLM’s DNA unwinding activity (Figure 6B) through the annealing of RPA-coated ssDNA bubbles. This SMARCAL1 activity was not observed when using the HD mutant, and was partially impaired when using a RPA-binding defective mutant (1N_1-30_) (Figure 6B). Moreover, ZRANB3 was unable to counteract BLM’s activity (Figure 6B), reinforcing the unique antagonistic functional relationship between BLM and SMARCAL1, and the ability of SMARCAL1, but not ZRANB3, to anneal RPA-coated ssDNA.

Consistent with a functional antagonism between SMARCAL1 and BLM, loss of SMARCAL1 increased BLM accumulation at telomeres, whereas exogenous SMARCAL1 expression suppressed telomeric BLM foci (Figure 6C). Both the catalytic activity of SMARCAL1 and its RPA-binding ability contributed to limiting BLM recruitment, as evidenced by the partial effects of the HD and ΔN_1-115_ mutants, indicating that multiple functional activities of SMARCAL1 are required to restrain BLM at ALT telomeres (Figure 6C). Conversely, overexpression of wild-type BLM, but not the catalytically inactive mutant K695A (KA) (Min et al. 2019), significantly enhanced SMARCAL1 localization at ALT telomeres (Figure 6D).

To directly assess the requirement for BLM helicase activity in mediating the phenotypes observed upon SMARCAL1 loss, endogenous BLM was depleted using a siRNA targeting the 3′UTR of the *BLM* mRNA, followed by complementation with exogenous BLM cDNA (Supplementary Figure 7A-G). Despite achieving only modest levels of nuclear protein expression, complementation with WT BLM, but not the KA mutant, partially restored SMARCAL1/TRF2 colocalization, DDR activation, APB and TIF formation, as well as the cell fitness impairment observed upon SMARCAL1 depletion (Supplementary Figure 7A-F). Importantly, consistent with a protective role for SMARCAL1 in counteracting BLM-helicase-driven telomeric stress, overexpression of WT BLM in control G292 cells did not induce appreciable telomeric RPA accumulation, APB formation or cell fitness impairment (Figure 6E-F, Supplementary Figure 7H). In contrast, overexpression of WT BLM, but not the KA mutant, markedly exacerbated DNA damage, telomeric RPA accumulation, APB formation and fitness defects in SMARCAL1-depleted cells (Figure 6E-F, Supplementary Figure 7G-H). Taken together, these findings suggest that BLM facilitates the formation of ssDNA intermediates at telomeres of ALT-positive cells, which in turn promote SMARCAL1 recruitment. By inhibiting PRIMPOL-mediated repriming and counteracting BLM-dependent DNA unwinding, SMARCAL1 could prevent excessive formation of ssDNA intermediates that would otherwise aggravate telomere instability and activate aberrant DDR signaling, resulting in an impairment of cancer cell fitness (Supplementary Figure 8G).

### SMARCAL1 loss induces senescence in ALT-positive cancer cells

Telomeric damage is a known driver of cellular senescence, a state of persistent growth arrest (Victorelli and Passos 2017; Barnes et al. 2022; De Rosa et al. 2023). Given the pronounced telomeric damage and cell fitness impairment observed upon SMARCAL1 depletion in ALT-positive cancer cells, we investigated whether these effects were accompanied by the induction of cellular senescence. Following SMARCAL1 depletion in OS384 and G292iDAXX cells, we detected an approximately five-fold increase in senescence-associated β-galactosidase (SA-β-gal)-positive cells (Figure 7A-B, Supplementary Figure 8A). Additional hallmarks of senescence, including morphological cellular changes (*e.g.*, increased nuclear area) and elevated *IL6* expression, were also observed in both G292iDAXX and SAOS2 cells (Supplementary Figure 8B-C). Notably, consistent with its effect in mitigating telomeric damage and restoring cell fitness, DAXX expression in G292iDAXX cells prevented the induction of senescence markers following SMARCAL1 depletion (Figure 7A-B, Supplementary Figure 8B).

**Figure 7.**
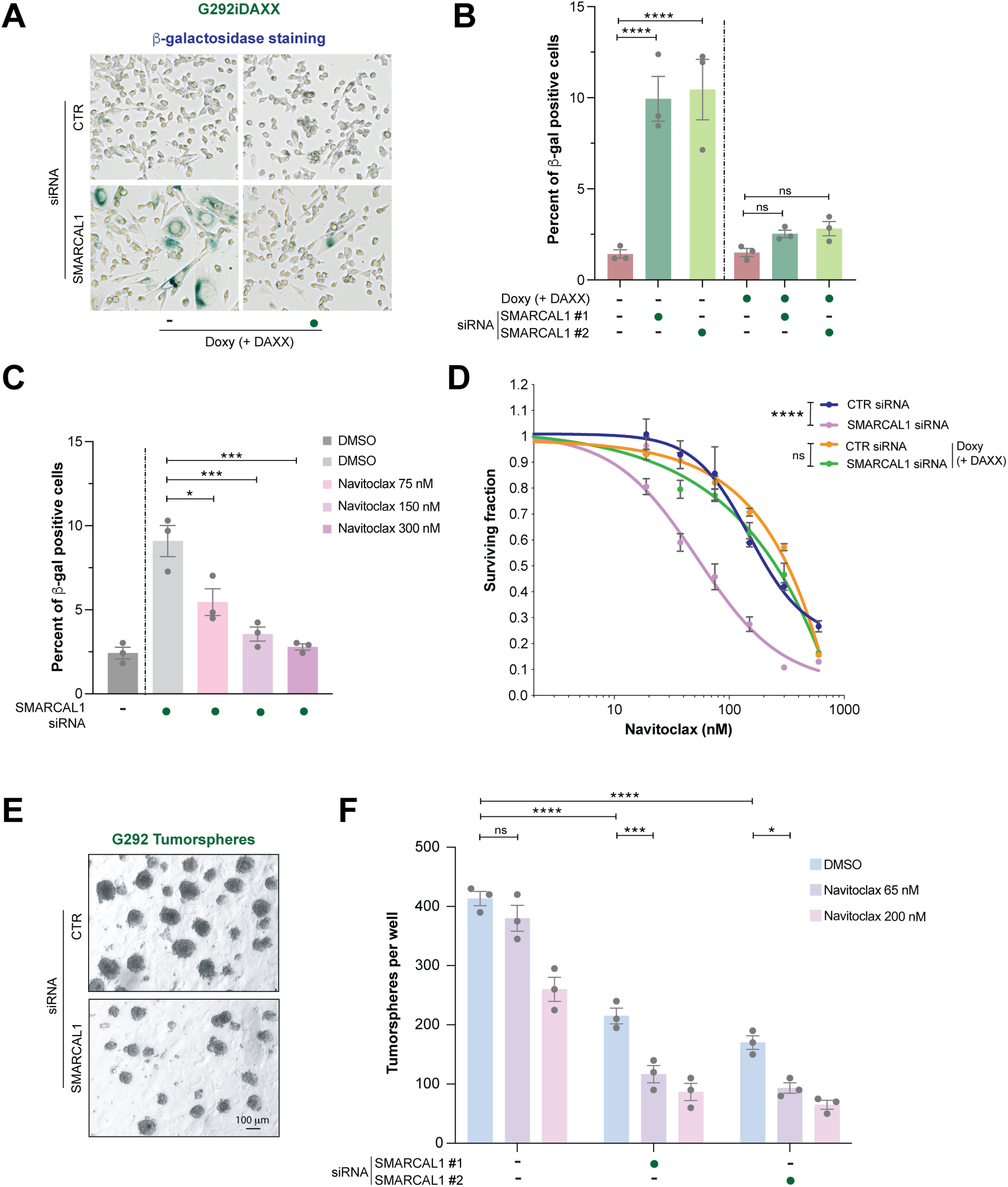
Senescence-associated phenotypes induced by SMARCAL1 loss in ALT-positive cancer cells **(A)** Representative images of β-galactosidase staining of G292iDAXX cells transfected with the indicated siRNAs, with or without doxycycline (doxy) treatment to express DAXX, as specified. **(B-C)** Quantification of the percentage of β-galactosidase (β-gal)-positive cells following transfection with the indicated siRNAs and treated as specified. Columns represent the mean ± SEM of independent biological replicates (dots). Statistical significance was determined using ordinary one-way ANOVA followed by Tukey’s multiple comparisons test. (D) Survival analysis in G292iDAXX cells following treatment with navitoclax at the indicated concentrations. Cell survival is expressed as a fraction relative to the untreated control, and data represent the mean ± SEM of at least three replicates per condition. Statistical significance was determined using ordinary two-way ANOVA followed by Tukey’s multiple comparisons test. **(E-F)** Representative images (E) and quantification (F) of G292 tumorspheres grown in non-adherent culture conditions following transfection with the indicated siRNAs and treated as specified. Columns represent the mean ± SD of independent biological replicates (dots), shown as the absolute number of spheres per well. Statistical significance was determined as in (B-C). Significance levels are denoted as follows: ns (not significant), *p < 0.05, **p < 0.01, ***p < 0.001, ****p < 0.0001.

We next hypothesized that the induction of cellular senescence upon SMARCAL1 depletion might sensitize ALT-positive cancer cells to senolytic agents, which selectively target senescent cells. To test this hypothesis, we examined the sensitivity of SMARCAL1-depleted, ALT-positive cancer cells to navitoclax, an inhibitor of anti-apoptotic BCL-2 family proteins (Tse et al. 2008). Increasing doses of navitoclax led to the selective clearance of senescent cells induced by SMARCAL1 depletion, confirming the specificity of the senolytic effect of navitoclax (Figure 7C). SMARCAL1-depleted ALT-positive cells showed marked sensitivity to navitoclax (Figure 7D, Supplementary Figure 8D) and to an additional senolytic agent, A1331852 (Zhu et al. 2017) (Supplementary Figure 8E). Importantly, DAXX induction, which suppressed cellular senescence, rendered SMARCAL1-depleted G292iDAXX cells resistant to senolytic agents (Figure 7D, Supplementary Figure 8E). In addition, depletion of either BLM or PRIMPOL rescued the increased sensitivity to navitoclax induced by SMARCAL1 loss in G292 cells (Supplementary Figure 8F). To further substantiate the therapeutic relevance of our findings, we performed tumorsphere growth assays under non-adherent culture conditions. SMARCAL1 depletion significantly impaired the growth of ALT-positive tumor spheroids, an effect that was further enhanced by treatment with low doses of navitoclax at which control tumorspheres are not responsive (Figure 7E-F). Together, these findings indicate that the fitness defects associated with SMARCAL1 depletion in ALT-positive cancer cells are accompanied by the induction of cellular senescence, a vulnerability that may be therapeutically exploited using senolytic agents to selectively target ALT-positive tumors (Supplementary Figure 8G).

## Discussion

The findings presented in this study highlight a critical role for the SNF2-family DNA translocase SMARCAL1 in maintaining telomere stability and cellular fitness in ALT-positive cancer cells. By leveraging genome-wide CRISPR screening data from the DepMap database, we show that *SMARCAL1* is the top dependency gene in telomerase-negative cancer cells (Figure 1), a finding independently corroborated by a recent study (Wu et al. 2025). In addition, our work establishes a causal relationship between SMARCAL1 and telomere stability that is specifically required for cell fitness under active ALT conditions. In particular, we observe that SMARCAL1 operates at telomeres in ALT-positive cells to limit DNA damage by suppressing ssDNA accumulation and APB formation (Figures 3-4, Supplementary Figures 3-4). These phenotypes are dependent on ALT activity, as demonstrated by their reversibility upon DAXX-mediated ALT inhibition in an isogenic DAXX-deficient osteosarcoma model (Figures 3-4, Supplementary Figure 3). The ability of SMARCAL1 to promote cell fitness and suppress telomeric RPA accumulation and APB formation in ALT-positive cells depends on both its RPA-binding and ATPase activities (Figure 3D-G, Supplementary Figure 3D-E), suggesting that SMARCAL1 may also exert ATP-independent functions at telomeres. These observations align with our prior biochemical studies showing that SMARCAL1 possesses a non-catalytic activity, dependent on its RPA-binding domain, that promotes annealing of RPA-coated single-stranded DNA (Halder et al., 2022).

Telomeric DNA damage and cell fitness defects are not observed upon loss of the SMARCAL1-related, SNF2-family DNA translocase ZRANB3 in ALT-positive cancer cells, in line with previous observations that ZRANB3 does not play a major function at telomeres (Poole and Cortez 2017) (Supplementary Figure 3G-H). Instead, increased telomeric damage and ALT activity has been observed in U2OS cells upon loss of the SNF2-family DNA translocase HLTF (Bai et al. 2024). While *ZRANB3* and *HLTF* have not been uncovered as differential dependency genes in ALT-positive cancer cells by DepMap analyses (Figure 1) (Lu and Pickett 2022; Salgado et al. 2024; Dubois et al. 2025; Wu et al. 2025), it remains to be determined whether their loss can exacerbate the dependency of ALT-positive cancer cells on SMARCAL1 and/or FANCM. Depletion of FANCM, an established suppressor of ALT activity in ALT-positive cancer cells, further increases telomere damage upon SMARCAL1 loss (Supplementary Figure 4C) (Lee et al. 2025). While our data suggest that telomere damage contributes to the synergistic loss of viability observed upon SMARCAL1 and FANCM co-depletion in ALT-positive cancer cells (Figure 2F, Supplementary Figure 2G-H), recent studies have also uncovered a strong synthetic lethal interaction between *SMARCAL1* and *FANCM* in non-transformed, ALT-negative epithelial cells (Feng et al. 2024; Agashe and Vindigni 2025; Fielden et al. 2025). These findings point to a redundant essential function of these enzymes independent of ALT that remains to be fully defined.

Our study also reveals that the previously described antagonistic interplay between SMARCAL1 and PRIMPOL (Quinet et al. 2020; Tirman et al. 2021) contributes to telomere stability in the context of ALT-positive cells (Supplementary Figure 8G). Specifically, we show that PRIMPOL overexpression exacerbates RPA phosphorylation and cell fitness defects caused by SMARCAL1 loss in ALT-positive cells (Figure 5A, Supplementary Figure 5A). As described in other cellular contexts (Piberger et al. 2020; Kang et al. 2021; Simoneau et al. 2021; Taglialatela et al. 2021; Tirman et al. 2021; Kawale et al. 2024; Pale et al. 2024), aberrant PRIMPOL repriming activity may favor accumulation of unrepaired telomeric ssDNA gaps and cellular toxicity. Functional analysis of PRIMPOL mutants supports this hypothesis, as overexpression of repriming and catalytically inactive, or RPA binding-deficient, PRIMPOL variants failed to exacerbate the cellular phenotypes induced by SMARCAL1 loss in ALT-positive cells (Figure 5B). However, the modest suppression of SMARCAL1-dependent phenotypes achieved by disrupting or depleting PRIMPOL (Figure 5C-E, Supplementary Figure 6) indicates that aberrant repriming alone cannot account for the full spectrum of defects induced by SMARCAL1 loss in ALT-positive cells.

Our observations that depletion of the BLM helicase fully suppresses the cellular effects induced by SMARCAL1 loss in ALT-positive cells (Figure 5E, Supplementary Figure 6) point to a critical interplay between BLM’s and SMARCAL1’s activities in ALT-positive cells (Supplementary Figure 8G). Accordingly, our findings support a direct role for SMARCAL1 in limiting BLM-mediated ssDNA accumulation and telomere instability (Figure 6, Supplementary Figures 7 and 8G). These results also emphasize BLM’s critical role in modulating telomere dynamics in ALT-positive cells. BLM’s DNA helicase activity, as part of the BTR complex, has been implicated in promoting ALT at multiple steps, including APB formation, replisome assembly, resolution of G-quadruplex DNA structures, and branch migration of DNA replication and recombination intermediates (Lu et al. 2019; Min et al. 2019; Loe et al. 2020; Zhang et al. 2021; O’Sullivan and Greenberg 2025). Recent studies suggest that BLM’s activity at telomeres promotes ALT by unwinding stalled Okazaki fragments on the G-rich strand, which arise from the stalling of the lagging-strand DNA polymerase at G-quadruplexes or other obstacles (Jiang et al. 2024; Lee et al. 2024). In these settings, BLM’s DNA helicase activity can generate C-rich 5’-ssDNA flaps, which may facilitate the recruitment of numerous DDR factors, promote APB formation, and enable telomere recombination and BIR events characteristic of ALT-positive cancer cells. Consistent with our findings (Figure 6D, Supplementary Figure 6A, D), proteomic analysis of telomeres in ALT-positive cells revealed that BLM loss abolishes the telomeric enrichment of multiple DDR factors, including SMARCAL1 (Jiang et al. 2024). However, SMARCAL1 is not enriched at telomeres in either ALT-positive or ALT-negative cells following induction of site-directed DNA double-strand breaks (DSBs), suggesting that its primary role is preventing telomeric replication stress rather than facilitating the repair of DSBs through BIR or other DSB repair pathways (Zhang et al. 2023). In support of a role for SMARCAL1 at difficult-to-replicate genomic regions, analysis of SMARCAL1-bound genomic sites revealed an enrichment for G-rich sequences prone to form G-quadruplex structures, raising the possibility that SMARCAL1 may operate at these sites (Leuzzi et al. 2024). Interestingly, HLTF has been recently shown to resolve G-quadruplexes (Bai et al. 2024), suggesting a function for SNF2-family DNA translocases in the metabolism of G-quadruplexes.

The above observations indicate that unscheduled BLM unwinding activity, licensed at telomeres of ALT-positive cells by frequent DNA polymerase stalling, may generate the DNA intermediates that trigger ALT. These same DNA structures could also serve as substrates for SMARCAL1, which in turn opposes BLM to restrain ALT (Supplementary Figure 8G). This hypothesis is supported by our *in vitro* biochemical studies demonstrating that SMARCAL1’s DNA annealing activity counteracts BLM’s DNA helicase activity (Figure 6A-B). The specific nature of the DNA structures generated by BLM that require SMARCAL1-dependent processing remains to be determined. Nonetheless, it is tempting to speculate a scenario in which SMARCAL1’s DNA remodeling activity extends beyond promoting fork reversal at stalled replication forks to include also post-replicative processes, such as limiting aberrant processing of ssDNA gaps or, as recently proposed, resolving R-loop intermediates (Hodson et al. 2022). Moreover, in the absence of SMARCAL1, BLM may contribute to the formation of ssDNA gaps at telomeres by cooperating with PRIMPOL to reprime DNA synthesis at stalled forks, as previously proposed in the context of interstrand crosslinks (Gonzalez-Acosta et al. 2021).

The significant induction of cellular senescence upon SMARCAL1 depletion highlights its critical role in maintaining telomere stability in ALT-positive cancer cells (Figure 7, Supplementary Figure 8). Our findings demonstrate that senescent ALT-positive cells induced upon SMARCAL1 loss are selectively sensitive to senolytic agents, such as navitoclax. This senescence phenotype was rescued by DAXX-mediated ALT suppression and loss of BLM or PRIMPOL, directly linking it to the telomere dysfunction caused by SMARCAL1 depletion (Figure 7, Supplementary Figure 8). These results suggest that the telomere stress generated upon SMARCAL1 loss may contribute to activate the DNA damage checkpoint response, ultimately leading to cell cycle arrest and senescence (Supplementary Figure 8G).

ALT activity in telomerase-negative cancer cells is governed by a complex network of independent and/or interrelated factors that must finely balance telomeric stress to allow BIR-mediated telomere elongation without compromising cell viability or proliferation. In most cases, ALT-positive tumors arise following the loss of ALT repressors, such as ATRX and DAXX. ATRX or DAXX loss, although not sufficient *per se* to establish a chronically active ALT phenotype (Lovejoy et al. 2012; O’Sullivan et al. 2014; Li et al. 2019; Turkalo et al. 2023), represents the most common route enabling ALT development during tumor evolution (Heaphy et al. 2011; Clatterbuck Soper and Meltzer 2023). Less commonly, ALT-positive tumors arise from hyperactivation of ALT inducers, such as TOP3A (de Nonneville et al. 2022). In these settings, SMARCAL1 may act as a “brake” to restrain ALT activity and maintain tolerable levels of telomeric stress. Consistent with this concept, *SMARCAL1* loss has been implicated in some tumors as the genetic alteration enabling the acquisition of ALT-positive phenotypes (Diplas et al. 2018; Brosnan-Cashman et al. 2021; Oak et al. 2025). Such tumors would not be expected to benefit from therapeutic strategies targeting SMARCAL1. However, our data (Figure 2D, F, Supplementary Figure 2E-J) suggest that therapeutically unleashing ALT by targeting FANCM or the ATRX/DAXX complex in tumors harboring SMARCAL1 loss may represent an alternative and potentially exploitable vulnerability. Indeed, loss of ATRX in *SMARCAL1*-mutant ALT-positive tumors is exceedingly rare, likely reflecting a requirement to finely balance ALT activity to sustain telomere elongation, cellular immortalization, and tumor growth (Diplas et al. 2018; Brosnan-Cashman et al. 2021; Liu et al. 2023). Importantly, a recent study identified SMARCAL1 as essential in ATRX-deficient, ALT-positive gliomas (Brown et al. 2026), further underscoring the synthetic lethal relationship between SMARCAL1 and ATRX loss in the context of ALT.

In conclusion, our study identifies SMARCAL1 as a pivotal regulator of telomere stability in ALT-positive cancer cells, revealing its interplay with BLM and PRIMPOL in maintaining ALT metabolism and cell fitness (Supplementary Figure 8G). The critical, selective and non-redundant function of SMARCAL1 in ALT-positive cancer cells highlights its potential as a promising therapeutic target. Moreover, the induction of senescence and the sensitivity of SMARCAL1-depleted cells to senolytic agents present additional avenues for clinical application (Supplementary Figure 8G). Future research aimed at developing potent and selective small-molecule inhibitors of SMARCAL1 and assessing their efficacy as standalone or combination therapies in preclinical models of ALT-positive cancers will be instrumental in advancing innovative treatment strategies targeting this distinctive mechanism of telomere maintenance.

## Materials and Methods

### Cell lines

MG63, HOS, SAOS2, HUO9 cancer cells were grown in DMEM with 10% FBS. G292 and G292iDAXX cells were grown in McCoy’s 5a Medium supplemented with 15% FBS. TM31 cells were grown in MEM with 10% FBS. SKLU1 cells were grown in Eagle’s Minimum Essential Medium with 10% FBS. PDX osteosarcoma #11890 cells were grown in SmGM-2 Smooth Muscle Cell Growth Medium supplemented with BulletKitL Lonza (cat. CC-3182) and 2 μl/ml of 5 μM Y-27632. #13063 cells were grown in MSCGM Mesenchymal Stem Cell Growth Medium supplemented with BulletKitT Lonza (cat. PT-3001) and 2 μl/ml of 5 μM Y-27632. #11721 cells were grown in 90% DMEM/F12 (1:1), 10% FBS, 1% NEAA, 1% L-glutamine, 200 mM; 2 μl/ml of 5 μM Y-27632. #86347 cells were grown in AR-5 complete growth medium (recipe available on request) with 2 μl/ml of 5 μM Y-27632. All cell lines were cultured in the presence of 100 U/ml of penicillin and 100 μg/ml of streptomycin at 37°C, 5% CO_2_.

### Plasmids

The doxycycline-inducible lentiviral vector for the expression of SpCas9 was previously described (Taglialatela et al. 2021). The gRNA targeting *SMARCAL1* (EX5_GTCTGCCTCGAAGTAGGCCC) used with inducible SpCas9 was cloned into pLenti-SpBsmBI (plasmid #62205, Addgene). The lentiviral plasmid for expression of enAsCas12a (pRDA_174) was obtained from Addgene (Plasmid #136476). Guide sequences were selected with CRISPick and shortened to 20 bp by 3 nucleotides from the 3’ ends. In4mer arrays were ordered as individual eBlocks (IDT) and cloned into the lentiviral Cas12a gRNA expression backbone pRDA_052 (Addgene #136474) using the restriction cut site Esp3I. The lentiviral expression vector pHAGE-Ct-Flag-HA-DEST-Puro to express siRNA resistant WT and mutant SMARCAL1 were previously described (Leuzzi et al. 2024). The SMARCAL1 catalytically inactive HD (D549A, E550A) mutant was generated by mutagenesis on the same backbone (Ciccia et al. 2012). Plasmids expressing V5-tagged WT PRIMPOL and mutants were generously provided by Juan Mendez (Mouron et al. 2013; Gonzalez-Acosta et al. 2021). pcDNA plasmids expressing GFP-tagged BLM were generously provided by Jerry W. Shay (Min et al. 2019).

### Recombinant viral production and cell transduction

Recombinant lentiviruses were generated by co-transfecting helper packaging vectors together with lentiviral vectors into HEK293T cells using the TransIT-293 transfection reagent (Mirus). Virus-containing supernatants were collected 48 hr after transfection, filtered and utilized to infect target cells in the presence of 8 μg/ml of polybrene. 48 hours after viral addition, transduced cells were selected using 1 μg/ml of puromycin, 100 μg/ml of hygromycin or 10 μg/ml of blasticidin for 3-5 days or until selection was successfully completed.

### Cell transfection

RNA interference experiments were carried out by reverse transfection of the indicated siRNAs using RNAiMAX reagents (#13778075, Thermo Fisher Scientific) according to the manufacturer’s instructions. For transfections involving DNA plasmids, 500 ng of DNA were combined with Lipofectamine 3000 (#L3000015, Thermo Fisher Scientific) according to the manufacturer’s instructions, and used for transfection of adherent cells at 50-60% confluency in 6-multiwell tissue culture plates.

### Generation of pooled KO and reconstituted cell lines

Doxycycline-inducible SpCas9-expressing G292 cells were obtained by lentiviral infection and blasticidin selection. Doxycycline-inducible SpCas9-expressing cells were further infected with lentiviral gRNA constructs targeting SMARCAL1 and selected with hygromycin. Cas12a-expressing cells were obtained by lentiviral infection and blasticidin selection. Cas12a-expressing cells were further infected with lentiviral In4mer gRNA constructs (Esmaeili Anvar et al. 2024) targeting the indicated genes and selected with puromycin. After 4-5 days of selection and recovery, cells were collected for experimental procedures and analysis of KO efficiency by WB and RT-qPCR.

### Cell fitness and survival assay

Transfected or transduced cells were seeded into 6-well plates at the same cell number (20-30% density) and expanded for 12-15 days. After approximately 8 doublings of control cells (split 1:4 at confluency of control cells for 3 times), cell fitness was determined by either counting viable cells using trypan blue, or by crystal violet staining and quantification. For crystal violet assays, cells were fixed and stained with a solution containing 1% formaldehyde and 1% crystal violet in methanol. The absorbed dye was resolubilized with methanol containing 0.1% SDS, which was then transferred into 96-well plates and measured photometrically (595 nm) in a microplate reader. Following background subtraction, cell viability was calculated by normalizing the absorbance of each sample to the control sample. For counting cells, the absolute cell number for each experimental condition was determined by using a hemocytometer counting chamber and then normalized to the relative control condition. For survival assays upon drug treatment, cells were seeded into 12-well plates at 10-20% density. Twenty-four hours after seeding, the cells were treated or not with the chemical agents described in the main text at the indicated concentrations. After incubation for 5-8 days, cell viability was assessed by crystal violet staining. Survival curves were generated using the nonlinear regression algorithm of the GraphPad Prism software.

### Tumorsphere formation assay

G292 cells were transfected with control and SMARCAL1 siRNAs on day 0. Twenty-four hours after transfection, cells were harvested and resuspended in complete culture medium supplemented with methylcellulose to a final concentration of 0.8%. Cells were seeded at a density of 10,000-15,000 cells per well in a final volume of 1 ml using 24-well low-attachment plates. Cultures were maintained without treatment for 5 days to allow tumorsphere formation. At this time point, 100 µl of fresh culture medium containing the indicated drug was added to each well to achieve the desired final concentration. Tumorspheres were then allowed to grow for an additional 5-7 days. For sphere quantification, cultures were collected into 15-ml conical tubes, brought to a final volume of 12 ml with PBS, and centrifuged at 600 rpm for 5 min. The supernatant was carefully removed by decanting or gentle aspiration, leaving 100-200 µl of residual PBS. Tumorspheres were resuspended, and the total number of spheres was determined by counting all spheres present in a 10-µl aliquot using a hemocytometer. Absolute sphere numbers per well were calculated by normalizing to the total resuspension volume.

### β-galactosidase senescence assay

After transfection, cells were cultured in 6-multiwell plates for approximately 6 doublings of control cells, then processed for β-galactosidase staining. The β-galactosidase cell staining (Cell Signaling #9860) was performed according to the manufacturer’s instructions. After staining, plates were imaged using an Evos microscope and the percentage of β-galactosidase positive senescent cells was determined by counting at least 3 images for each experimental condition (n of cells > 150). The CellEvent Senescence Green Detection Kit (Invitrogen #C10851) was used to determine β-galactosidase positive cells by fluorescence and automated imaging. The percentage of β-galactosidase positive senescent cells was determined by counting scoring > 1000 cells for each experimental condition.

### Western blotting

Cells were collected by trypsinization and centrifugation (1500 x g at 4°C for 5 min). Cell pellets were washed in PBS twice, counted and resuspended in lysis buffer (37.5 mM tris-HCl pH 6.8, 1.25% SDS, 10% glycerol, 3% β-mercaptoethanol, 0.002% bromophenol blue), sonicated on ice and boiled for 5 min at 95°C. Sonication was performed on the Fisher Scientific Sonic Dismembrator Model 100 instrument (low power output, level 7, 10 sec ON / 60 sec OFF double pulses). Protein lysates were resolved on a 4-15% Mini-Protean TGX precast polyacrylamide gels (#456-108, Bio-Rad) and transferred to 0.45 μm nitrocellulose membrane (#HATF00010, Millipore). Membranes were incubated for 1 hr in TBS-T (50 mM tris-HCl pH 8, 150 mM NaCl, 0.1% Tween-20) containing 5% non-fat dry milk and then incubated for 2 hr at room temperature (RT) or overnight at 4°C with primary antibodies. Detection was achieved using appropriate horseradish peroxidase (HRP)-conjugated secondary antibodies.

### RNA extraction and RT-qPCR assay

All reagents, buffers and containers involving RNA work were RNase-free grade. Total RNA was isolated using the RNeasy kit (#74104, Qiagen) according to the manufacturer’s instructions. For real-time quantitative PCR (RT-qPCR) assays, 1 μg of total RNA was reverse-transcribed into cDNA using random hexamer primers (#N8080127, Thermo Fisher Scientific) and MMLV High Performance Reverse Transcriptase kit (#RT80125K, Lucigen). An equal amount of cDNA from different samples was mixed with Power SYBR green PCR master mix (#4367659, Thermo Fisher Scientific). Gene-specific primers were used for RT-qPCR amplification performed on a QuantStudio 3 System instrument. Gene-specific mRNA level for each experimental condition was determined using the ΔΔCT method and normalized to *TBP* mRNA. RT-qPCR results were presented as fold change of gene expression in the test samples compared to the control.

### Immunofluorescence and FISH

Cells were seeded in black 96-multiwell bottom-glass plates or grown on coverslips in 6-multiwell plates. When culture plates were 80-90% confluent (2-3 days after seeding), cells were simultaneously fixed and permeabilized (4% paraformaldehyde, 0.5% Triton X-100) for 10 min at RT. Fixed cells were incubated in blocking solution (3% BSA in TBS-Tween 0.1%) for 1 hr and then incubated for 1.5 hr at RT or overnight at 4°C with the primary antibodies. Cells were washed 3 times with TBS-T and incubated for 1 hr at RT with the appropriate labeled secondary antibody. After three washes in TBS-T, cells were incubated with DAPI for 5 min at RT to counterstain nuclei.

Telomere FISH was performed using fluorescently labeled telomere probes (PNA bio) according to the manufacturer’s instructions. For IF-FISH, following the IF protocol, coverslips were denatured and hybridized with the PNA probe (PNA Bio). Multi-color acquisitions were made using the ImageXpress Nano Automated Imaging System microscope or the Nikon Eclipse 50i microscope equipped with a 40× Plan Apo objective (0.95 numerical aperture). MetaXpress6 software was used for automated image analysis.

### Native telomere FISH

Native telomere FISH was performed as previously described (Loe et al. 2020; Azeroglu et al. 2024). Cells were grown on coverslips and fixed in 2% paraformaldehyde for 5 min. Coverslips were then incubated with 500 µg/ml of RNase A in blocking solution (1 mg/ml BSA, 3% goat serum, 0.1% Triton X-100, 1 mM EDTA in PBS) for 1 hr at 37°C. Coverslips were dehydrated in an ethanol series (70%, 90%, and 100%) and hybridized at room temperature with a Cy3-OO[TTAGGG]3-labeled PNA probe (#F1006, PNA Bio) in hybridization buffer (70% formamide, 1 mg/ml blocking reagent [Roche], 10 mM tris-HCl at pH 7.2). Following washes, coverslips were stained with DAPI and imaged. Single-plane images were acquired using a Zeiss Axio Imager M2 with an Axiocam 702 camera and a Zeiss Axio observer with an Axiocam 712 camera using ZEN 2.6 (blue edition) software.

### 4SET assay

The 4SET assay was performed as previously described (Lee et al. 2024). Briefly, cells were harvested using trypsin-EDTA and then divided into separate tubes for fractionation. DNA was purified from the soluble fractions, and 150 ng of soluble DNA (quantified using nanodrop and Qubit) was loaded into a 0.5% agarose gel in TAE buffer (Tris-acetate EDTA). The native agarose gel was transferred onto a positively charged Nylon membrane (Roche) and subjected to Southern blot with (Digoxigenin)-labeled strand specific telomere probes. The membrane was incubated with 1% CDP-star (Roche 11759051001) in AP buffer (50 mM Tris pH 9.5, 100 mM NaCl) for signal detection, which was performed using the Odyssey Imaging System (LI-COR 2800).

### Topo I-coupled unwinding of covalently closed plasmid DNA

Reactions (20 μl) containing 25 mM tris-acetate pH 7.5, 1.8 mM magnesium acetate, 20 mM NaCl, 1 mM dithiothreitol, 100 μg/ml recombinant albumin, 2 mM ATP, 5 ng/μl pUC19 negative scDNA were assembled on ice. pUC19 was relaxed by incubation with 16.5 nM *Drosophila* Topo I for 10 min at 37°C. DNA unwinding reactions were then initiated by adding 47 nM BLM in the presence of 936 nM RPA and incubating for 5 min at 37°C. The reactions were subsequently supplemented with the indicated concentration of SMARCAL1 variants and ZRANB3. After incubation for 25 min, the reactions were terminated by introducing NaCl to a final concentration of 555 mM for 2 min at RT, followed by the addition of 8 µl of STOP buffer (150 mM EDTA, 2% SDS, 30% glycerol and 0.1% bromophenol blue) and 2 µl proteinase K (14-22 mg/ml; Roche) for 20 min at 37 °C. Reaction products were analyzed by electrophoresis (4 V/cm for 90 min) in 1% agarose-TAE buffer (40 mM tris-acetate pH 8.0, 1 mM EDTA). DNA was stained with TAE buffer containing GelRed (1:10,000, Biotium) for 30 min with gentle agitation after the completion of electrophoresis. Images of gels were acquired with Quantum (Vilber). Results were quantified using ImageJ and plotted using Prism software (Prism 10, GraphPad).

### Protein purification

Recombinant wild-type FLAG-SMARCAL1, its catalytically inactive (HD; D549A, E550A) and RPA binding-deficient (DN_1-30_, lacking the first 30 amino acids) variants were expressed using pFastBac-FLAG-SMARCAL1 expression constructs in insect *Spodoptera frugiperda* 9 (*Sf*9) cells, and purified by affinity chromatography (Ciccia et al. 2012). N-terminally truncated *Drosophila* topoisomerase I (Topo I, catalytic subunit) containing a 6x His-tag at the C-terminus was expressed in *E. coli.* The cells were lysed by sonication. Topo I was purified using Ni-NTA resin and then applied on HiTrap S and HiTrap Heparin columns connected in tandem (GE Healthcare), and eluted from the HiTrap Heparin column after the HiTrap S column was disconnected (Halder et al. 2022). Human RPA was purified by using ion exchange and affinity chromatography (Ceppi et al. 2024). BLM was purified using the maltose-binding protein (MBP) tag at the N-terminus and 10x His-tag at the C-terminus as described in (Pinto et al. 2016).

### Differential gene dependency analysis

The gene effect dataset of CRISPR-Cas9 essentiality screens in 1,009 pan-cancer cell lines (gene effect scores derived from CRISPR knockout screens published by Broad’s Achilles and Sanger’s SCORE projects, release public 2023q4) (Meyers et al. 2017; Dempster et al. 2021; Pacini et al. 2021) and their gene expression profiles were downloaded from the Cancer Dependency Map portal (DepMap, https://depmap.org/portal/). Dependency score was the opposite number of the gene effect value for each gene, as described previously (Chen et al. 2023). Differential gene dependencies between cell line groups (telomerase-positive or -negative) was calculated using student’s *t*-test in R (parameters: alternative = ‘two.sided’, mu = 0, paired = FALSE, var.equal = FALSE, conf.level = 0.95) and the p values were adjusted using the R function p.adjust (parameters: method = ‘BH’). The differential gene dependencies were visualized using the R package ggplot2 (v3.2.1). We performed hierarchical clustering on gene dependencies of cell lines using the R package pheatmap (Pretty Heatmaps v1.0.10, parameters: clustering_method = ‘ward.D2’, clustering_distance_cols = ‘euclidean’, and cutree_cols = 2).

### Statistical analysis

Statistical analysis was performed using the GraphPad Prism software (v10.0), unless otherwise specified. Statistical analysis details for the different experiments are reported in figure legends or the methods section. In all cases: ns not significant; ∗ p<0.05; ∗∗ p<0.01; ∗∗∗ p<0.001; ∗∗∗∗ p<0.0001. Quantitative data are either presented as the mean ± SEM of at least 3 independent experiments or distribution of pooled data from three or more independent biological replicates. Images shown are representative of two or more independent experiments with similar results.

### Software used

Microsoft Excel, MetaXpress (v6), Rstudio, ImageJ and GraphPad Prism (v10.0) were utilized in this study.

## Competing interest statement

The authors declare no competing interests.

## Acknowledgments

The authors would like to thank Alison Taylor, Chao Lu and all members of the Ciccia laboratory for helpful suggestions and critical discussions, Sarah Clatterbuck Soper and Paul Meltzer for providing G292 and G292iDAXX cells (Yost et al. 2019), Juan Mendez for providing antibody for PRIMPOL detection and plasmids expressing V5-tagged WT and mutant PRIMPOL (Mouron et al. 2013; Gonzalez-Acosta et al. 2021), Shivi Dua for assistance in the generation of preliminary data. This work was supported by NIH grant R01CA197774 to A.C., R35GM155138 to J.M., the Intramural Research Program of the National Institutes of Health, National Cancer Institute and Center for Cancer Research, project 1-ZIA-BC011815-03 to E.L.D, and the Swiss Cancer Research grant KFS-6136-08-2024 to P.C..

## Author contributions

A.T. and A.C. conceived the study. A.T. and A.C. supervised all the work if not otherwise specified. A.T. performed the experimental work if not otherwise specified. A.G. and Z.K. helped A.T. to generate cell lines, perform cell fitness, survival, IF, FISH and western blotting assays. C.X. performed the analysis of the DepMap CRISPR screens. B.A. performed the native Tel-FISH experiments under the supervision of E.L.D.. J.L. performed the 4SET assays under the supervision of J.M.. M.R.D.S. performed the Topo I-coupled plasmid assay under the supervision of P.C.. G.L. provided siRNA resistant constructs for SMARCAL1 reconstitution. T.D. designed and cloned the In4mer gRNA constructs for Cas12a. A.V. provided viral particles for cell transduction and help with construct cloning. F.D. and A.K. provided osteosarcoma PDX derived cells. E.L.D. and J.L. provided reagents, expertise and feedback. A.T. and A.C. wrote the manuscript with contributions and input from all authors.

## Supplementary Figures

**Supplementary Figure 1.**
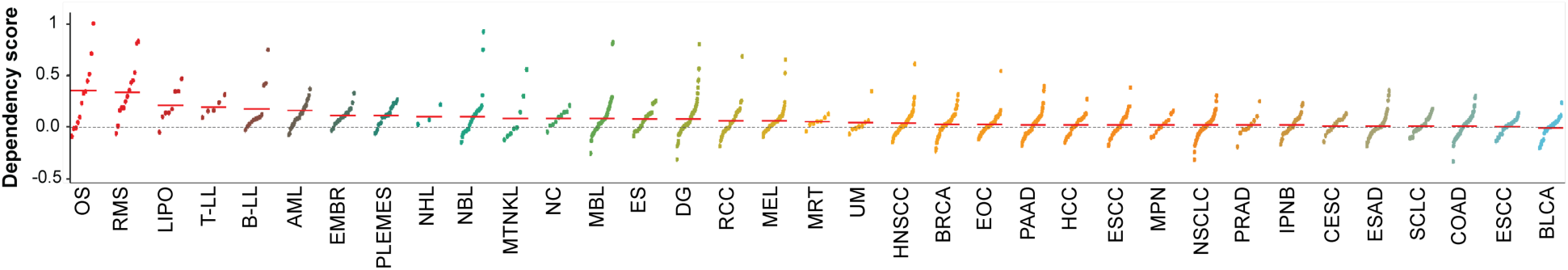
SMARCAL1 dependency score across cancer cell types Dot plot showing SMARCAL1 dependency in cancer cell lines grouped based on primary cancer disease type. Mean dependency scores are denoted by red bars. P-values were determined by unpaired two-tailed Student’s t-test.

**Supplementary Figure 2.**
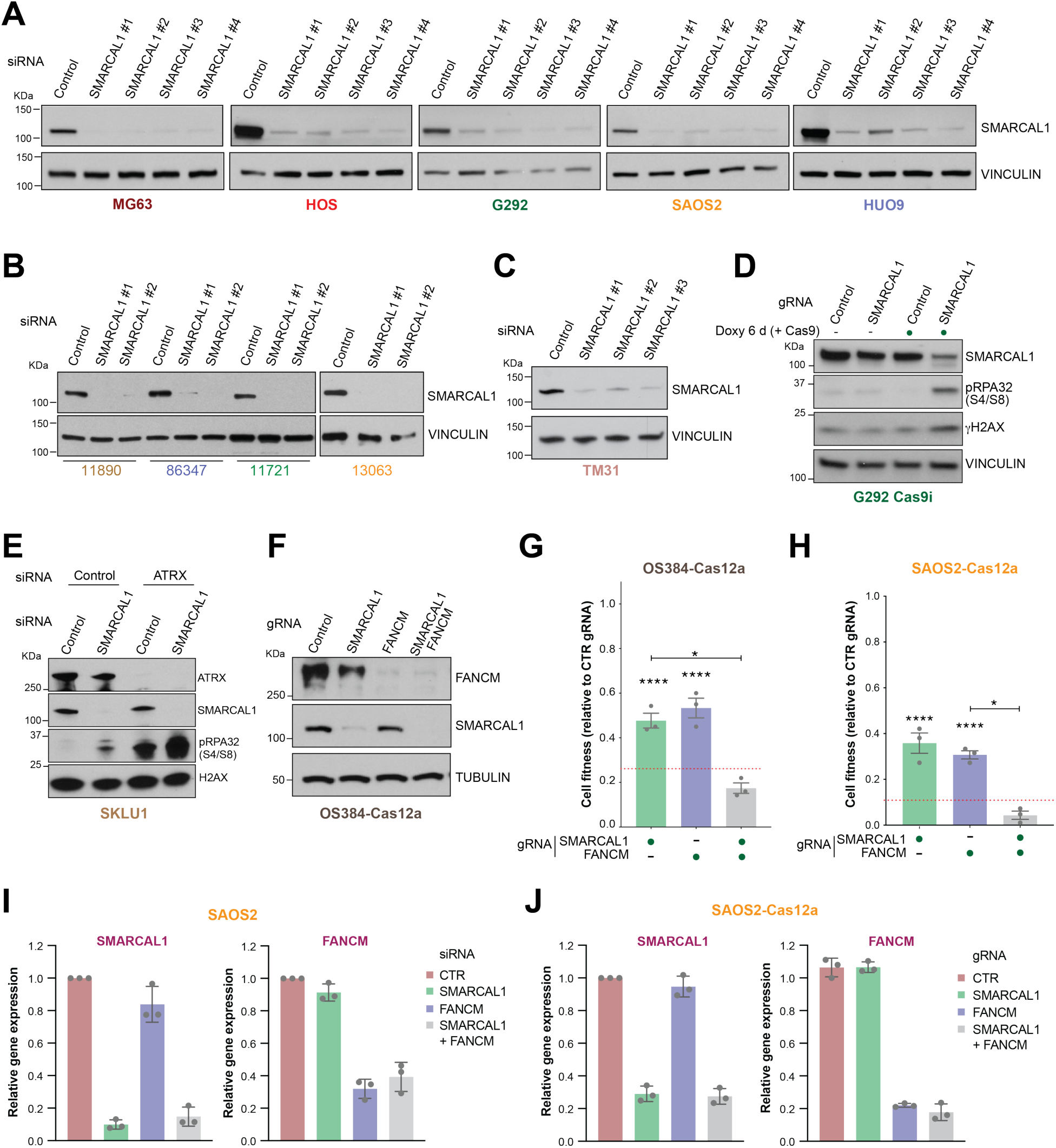
Effects of SMARCAL1 and/or FANCM loss in ALT-positive cancer cells **(A-F)** Detection by western blotting of SMARCAL1, phosho-RPA32 (S4/S8), γH2AX and FANCM levels in whole cell extracts of the specified cells treated as indicated. Vinculin, tubulin and H2AX are shown as a loading control. **(G-H)** Quantification of the fitness of OS384 (G) and SAOS2 (H) cells expressing Cas12a and transduced with the indicated gRNAs relative to the control gRNA. Columns represent the mean ± SEM of independent biological replicates (dots). The expected effect induced by dual gene targeting is indicated by the dashed red line. Statistical significance was determined using ordinary one-way ANOVA followed by Tukey’s multiple comparisons test. **(I-J)** Detection by RT-qPCR of the fold change in *SMARCAL1* and *FANCM* mRNA levels in SAOS2 cells treated with the indicated siRNAs (I) or expressing Cas12a and the indicated gRNAs (J) relative to the control treatment (CTR). *SMARCAL1* and *FANCM* mRNA levels in both cell lines were normalized to *TBP* mRNA levels. Significance levels are denoted as follows: *p < 0.05, ****p < 0.0001.

**Supplementary Figure 3.**
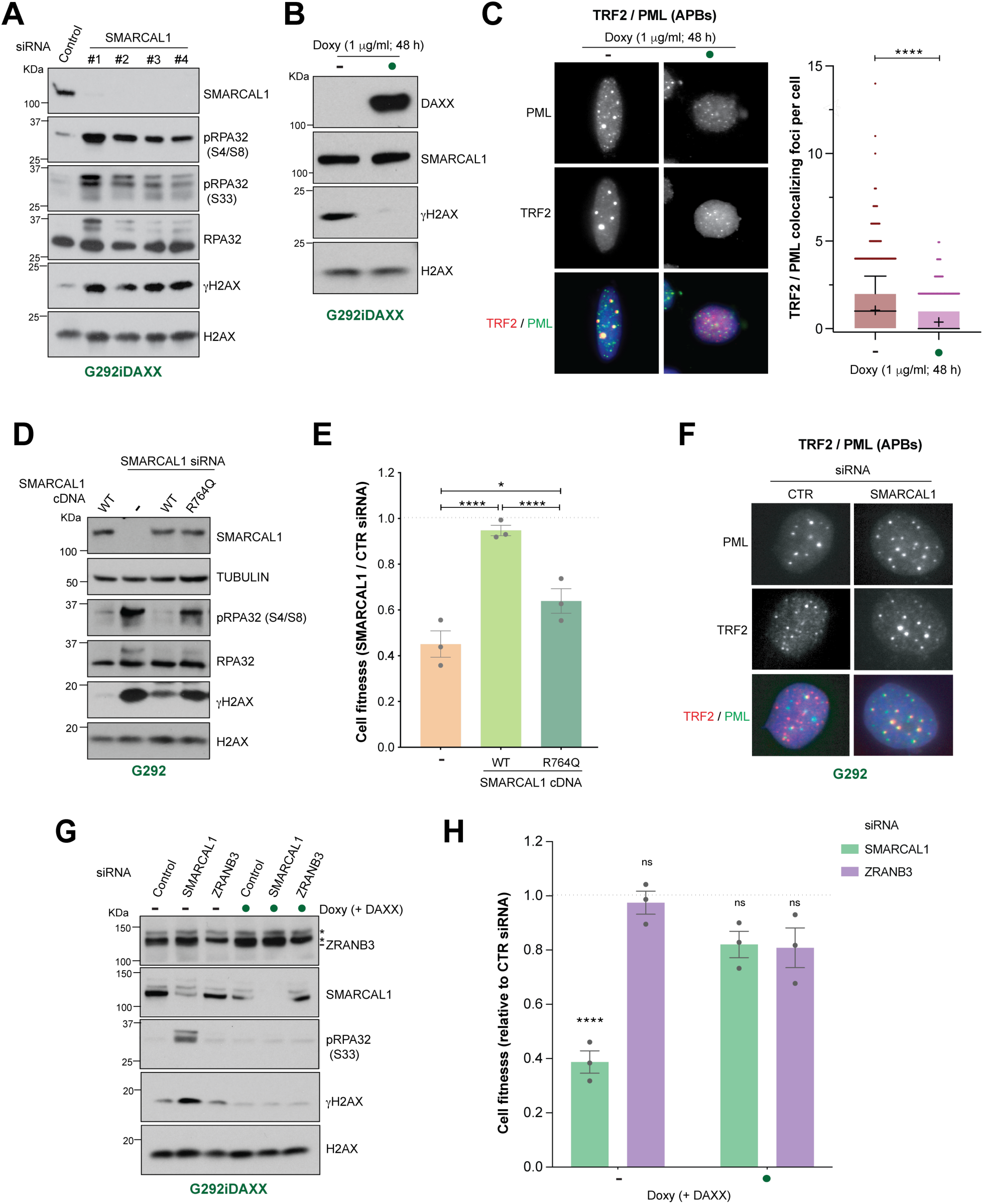
DNA damage signaling and cell fitness effects induced by the loss of SMARCAL1 or ZRANB3 in ALT-positive cancer cells **(A, B, D, G)** Detection by western blotting of the indicated proteins in whole cell extracts of the indicated cells, treated as specified. Total RPA32 and H2AX are included as loading controls. Asterisks indicate non-specific bands. **(C, F)** Representative images and quantification of PML foci per nucleus colocalizing with TRF2 in G292iDAXX (C) and G292 (F, related to Figure 3F) cells treated as indicated. Data are represented as box-and-whisker plots, showing the median (line), mean (+), 1^st^ to 3^rd^ quartiles (colored box), whiskers extending to the 10^th^ and 90^th^ percentiles, and individual outlier points. Data were obtained from 3 independent experiments. Statistical significance was determined using ordinary one-way ANOVA followed by Tukey’s multiple comparisons test. (E) Quantification of the fitness of SMARCAL1-depleted (siRNA) G292 cells relative to the control condition (CTR siRNA), treated as in (D) to express the indicated SMARCAL1 cDNA constructs. Columns represent the mean ± SEM of independent biological replicates (dots). Statistical analysis was conducted as in (C, F). **(H)** Quantification of the fitness of G292iDAXX cells transfected as in (G) with the indicated siRNAs relative to the control condition (CTR siRNA, dashed grey line), with or without doxycycline treatment. Graphical representation and statistical analysis were conducted as in (E). Significance levels are denoted as follows: ns (not significant), *p < 0.05, **p < 0.01, ***p < 0.001, ****p < 0.0001.

**Supplementary Figure 4.**
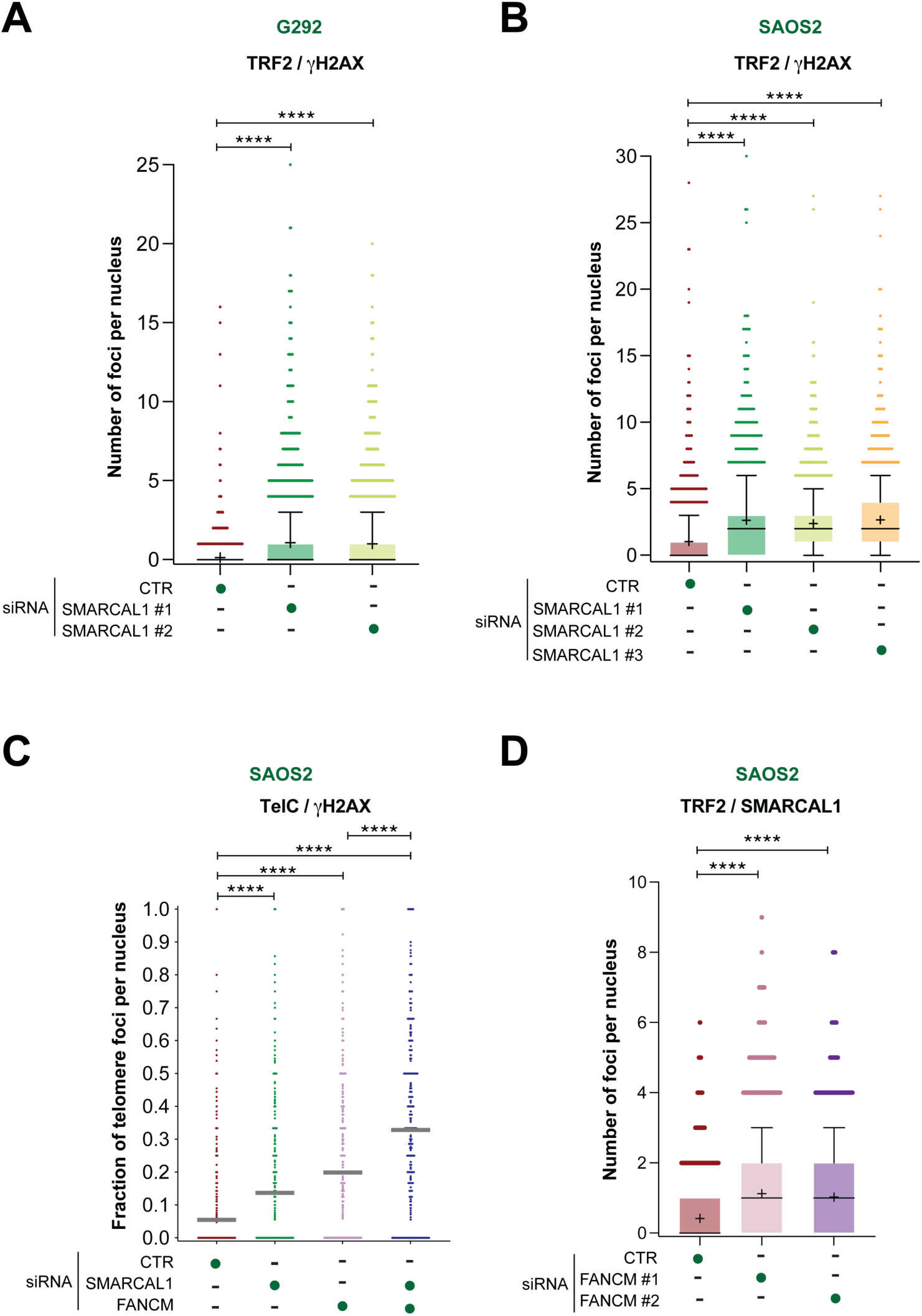
Telomeric DNA damage induced upon loss of SMARCAL1 and/or FANCM in ALT-positive cancer cells **(A-B)** Quantification of the number of γH2AX foci per nucleus colocalizing with TRF2 in G292 (A) and SAOS2 (B) cells transfected with the indicated siRNAs. Data are represented as box-and-whisker plots, showing the median (line), mean (+), 1^st^ to 3^rd^ quartiles (colored box), whiskers extending to the 10^th^ and 90^th^ percentiles, and individual outlier points. Data were obtained from 3 independent experiments. Statistical significance was determined using ordinary one-way ANOVA followed by Tukey’s multiple comparisons test. **(C)** Quantification of the fraction of telomeres positive for γH2AX per nucleus by IF-FISH. Data are represented as a dot plot, with the mean values indicated by grey lines. Data were derived from 3 independent experiments. Statistical analysis was conducted as in (A-B). **(D)** Quantification of the number of SMARCAL1 foci per nucleus colocalizing with TRF2 in SAOS2 cells transfected with the indicated siRNAs. Data representation and statistical analysis were conducted as in (A-B). Significance levels are denoted as follows: ns (not significant), *p < 0.05, **p < 0.01, ***p < 0.001, ****p < 0.0001.

**Supplementary Figure 5.**
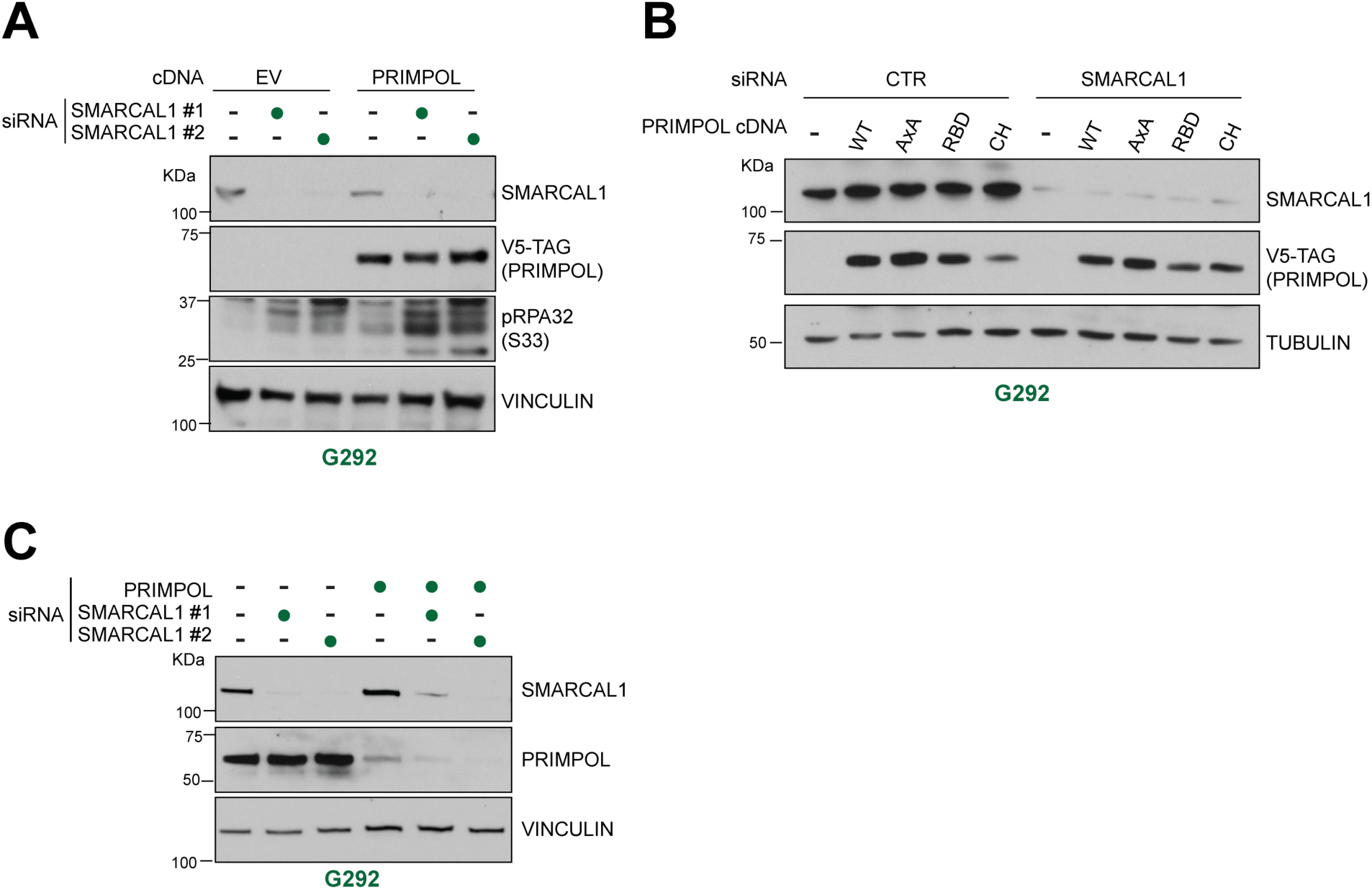
Protein levels following depletion of SMARCAL1 and/or PRIMPOL in ALT-positive cancer cells **(A-C)** Detection by western blotting of the indicated proteins in whole cell extracts of the specified cells treated with the indicated cDNAs or siRNAs. Vinculin and tubulin are shown as a loading control.

**Supplementary Figure 6.**
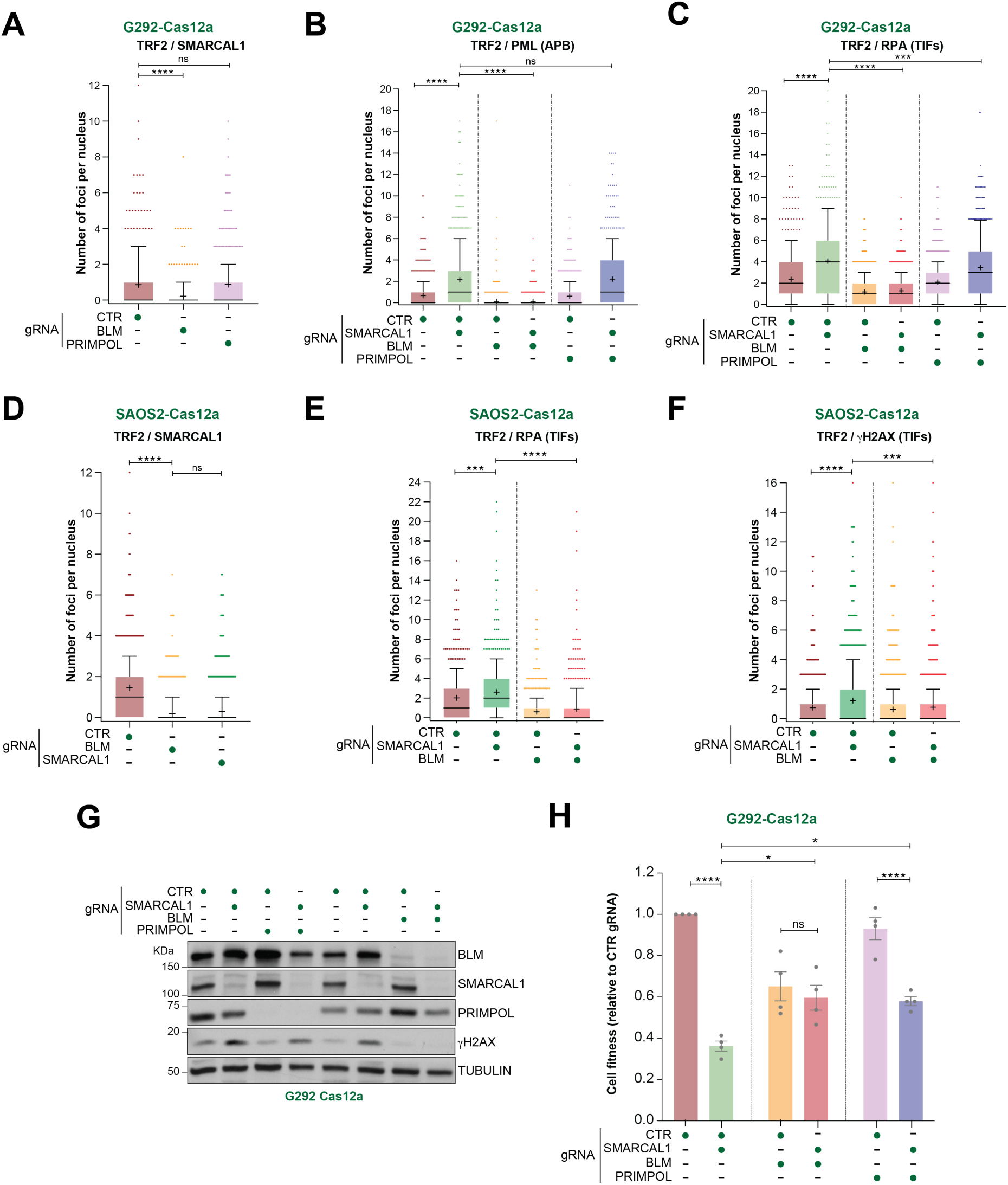
Telomeric foci and cell fitness in SMARCAL1-deficient ALT-positive cancer cells upon BLM or PRIMPOL loss **(A-F)** Quantification of the indicated colocalizing foci per nucleus in G292 (A-C) and SAOS2 (D-F) cells expressing Cas12a and transduced with the indicated gRNAs. Data are represented as box-and-whisker plots, showing the median (line), mean (+), 1^st^ to 3^rd^ quartiles (colored box), whiskers extending to the 10^th^ and 90^th^ percentiles, and individual outlier points. Data were obtained from 3 independent experiments. Statistical significance was determined using ordinary one-way ANOVA followed by Tukey’s multiple comparisons test. **(G)** Detection by western blotting of the indicated proteins in whole cell extracts of G292-Cas12a cells treated with the indicated gRNAs. Tubulin is shown as a loading control. **(H)** Quantification of the fitness of G292 cells expressing Cas12a and transduced with the indicated gRNAs, relative to the control conditions (CTR gRNA), as determined by cell counting. Graphical representation and statistical analysis were conducted as in (A-F). Columns represent the mean ± SEM of independent biological replicates (dots). Significance levels are denoted as follows: ns (not significant), *p < 0.05, **p < 0.01, ***p < 0.001, ****p < 0.0001.

**Supplementary Figure 7.**
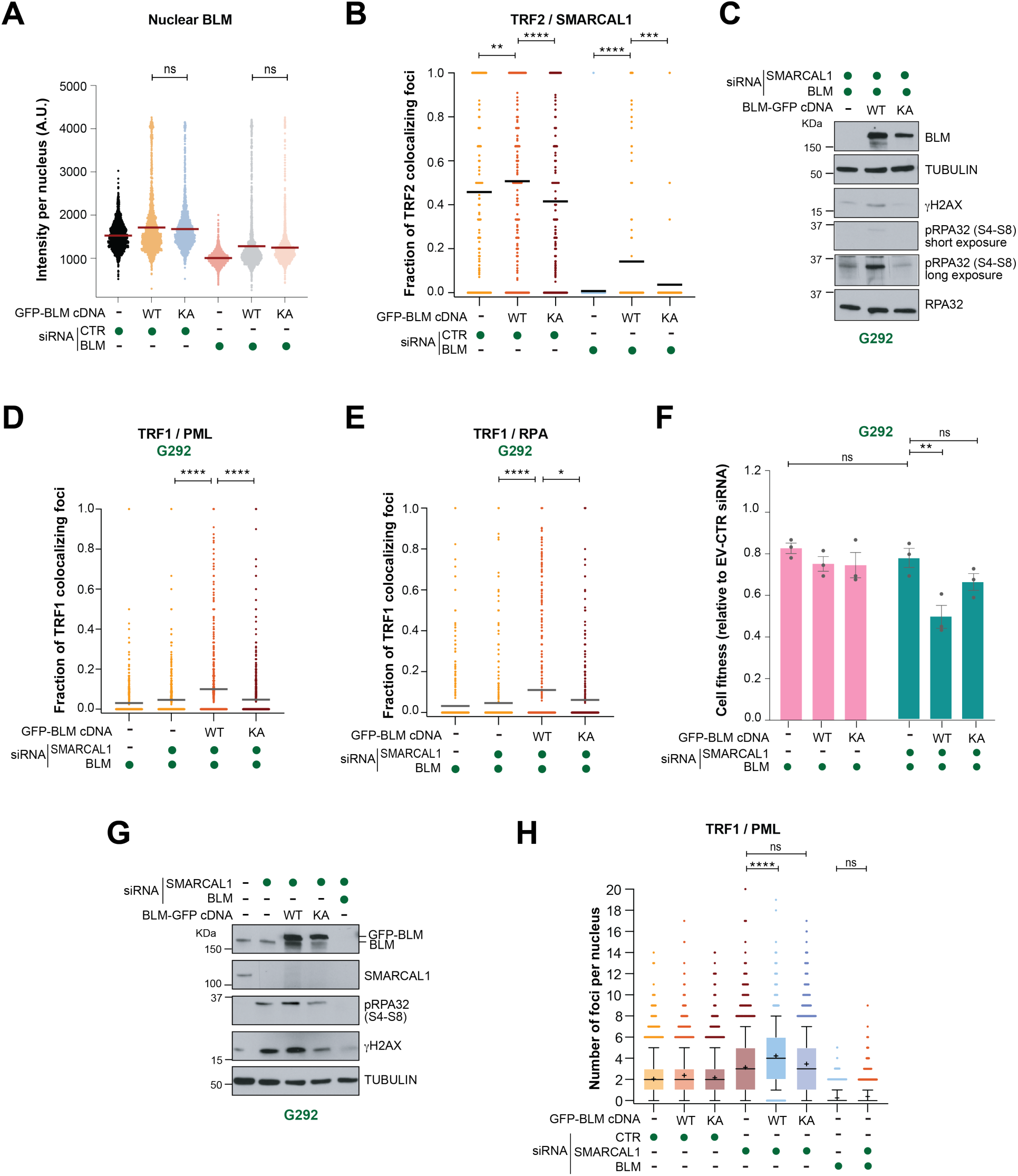
Cellular effects induced by BLM mutants in ALT-positive G292 cells **(A)** Nuclear BLM protein level upon transfection with the indicated cDNAs and siRNAs. BLM intensity per nucleus was detected by immunostaining and automated imaging of nuclear signal. Data are represented as a dot plot, with the mean values indicated by red lines. Statistical comparison was determined using ordinary one-way ANOVA followed by Tukey’s multiple comparisons test. **(B, D, E)** Quantification of the fraction of TRF2 foci positive for SMARCAL1 per nucleus (B), TRF1 foci positive for PML (D) or RPA32 (E) in G292 cells treated with the indicated cDNAs and siRNAs. Data are represented as a dot plot, with the mean values indicated by grey lines. Data were derived from 3 independent experiments. Statistical analysis was conducted as in (A). **(C, G)** Detection by western blotting of the indicated proteins in whole cell extracts of G292 cells treated with the indicated cDNA and siRNAs. Tubulin and RPA32 are shown as a loading control. **(F)** Quantification of the fitness of G292 cells by cell counting upon transfection with the indicated siRNAs and cDNAs, relative to cells treated with an empty vector (EV) and a control siRNA (CTR siRNA). Columns represent the mean ± SEM of independent biological replicates (dots). Statistical analysis was conducted as in (A). **(H)** Quantification of the indicated colocalizing foci per nucleus in G292 cells transfected with the indicated cDNA and siRNAs. Data are represented as box-and-whisker plots, showing the median (line), mean (+), 1st to 3rd quartiles (colored box), whiskers extending to the 10th and 90th percentiles, and individual outlier points. Data were obtained from 3 independent experiments. Statistical significance was determined as in (A). Significance levels are denoted as follows: ns (not significant), *p < 0.05, **p < 0.01, ***p < 0.001, ****p < 0.0001.

**Supplementary Figure 8.**
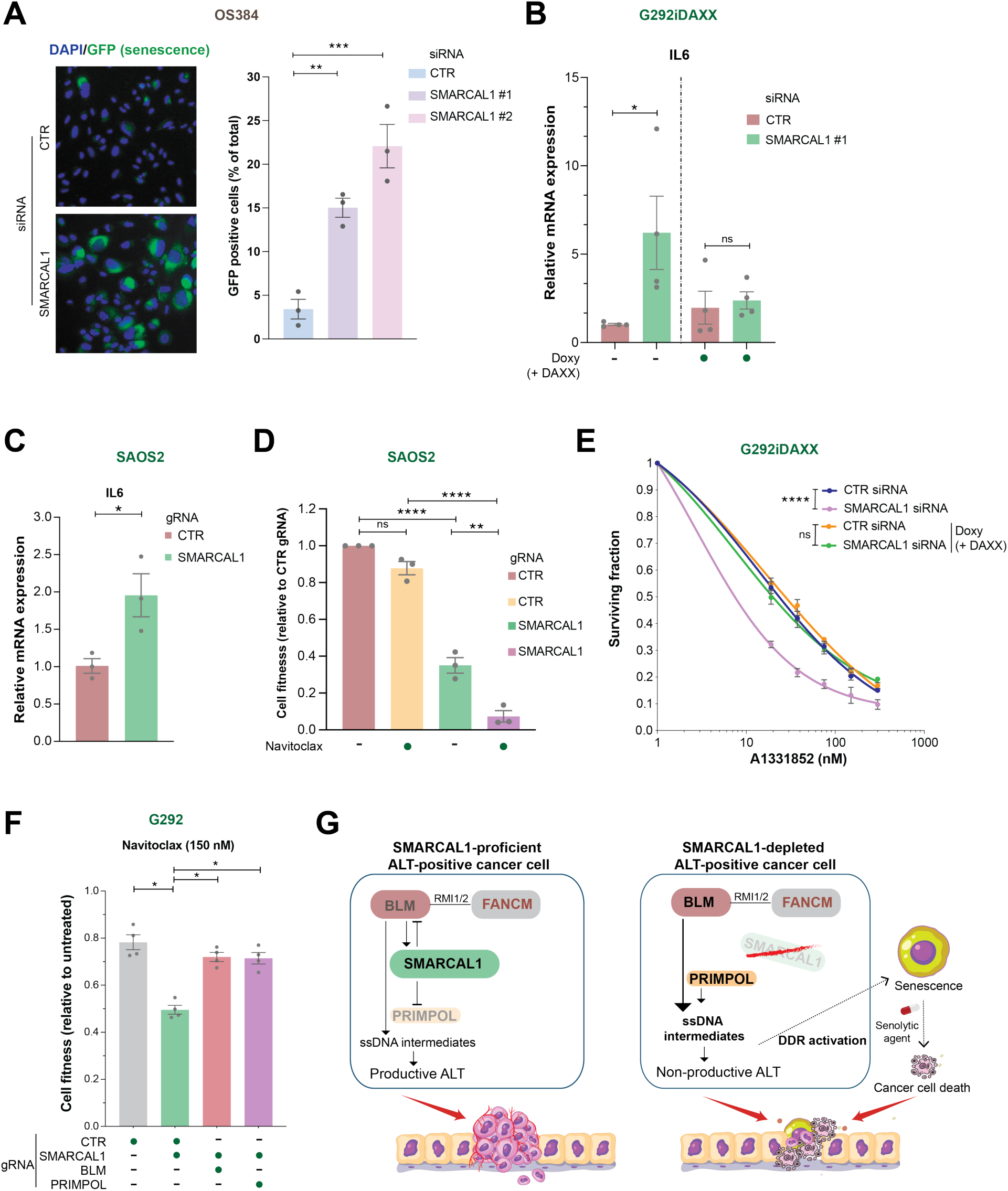
Senescence-associated phenotypes in SMARCAL1-deficient cells and schematic representation of the interplay between BLM, SMARCAL1 and PRIMPOL in ALT-positive cancer cells **(A)** Representative images of nuclei and β-galactosidase fluorescence in OS384 cells transfected with the indicated siRNAs (left panel). Quantification of the percentage of senescent (β-gal)-positive cells following transfection with the indicated siRNAs (right panel). Columns represent the mean ± SEM of independent biological replicates (dots). Statistical significance was determined using ordinary one-way ANOVA followed by Tukey’s multiple comparisons test. **(B-C)** Quantification of *IL6* mRNA levels by RT-qPCR in G292iDAXX (B) and SAOS2 cells expressing Cas12a (C) treated with the indicated siRNAs and gRNAs, respectively, with or without doxycycline (doxy) treatment to express DAXX, as indicated. *IL6* mRNA levels were normalized to *TBP* mRNA levels and shown as fold change relative to the control condition (CTR). Data representation and statistical analysis were conducted as in (A). **(D)** Quantification of the fitness of SAOS2 cells transduced with the indicated gRNAs, with or without navitoclax (0.3 mM) treatment, relative to the control condition (CTR gRNA). Columns represent the mean ± SEM of independent biological replicates (dots). Statistical significance was determined using ordinary two-way ANOVA followed by Tukey’s multiple comparisons test. **(E)** Survival of G292iDAXX cells following treatment with A1331852 at the indicated concentrations. Cell survival is expressed as a fraction relative to the untreated control, and data represent the mean ± SEM of at least three replicates per condition. Statistical analysis was conducted as in (A). **(F)** Quantification of the fitness of G292 cells expressing Cas12a and transduced with the indicated gRNAs, with navitoclax (150 nM) treatment relative to the untreated condition. Graphical representation and statistical analysis were conducted as in (A). Significance levels are denoted as follows: ns (not significant), *p < 0.05, **p < 0.01, ***p < 0.001, ****p < 0.0001. **(G)** Graphical schematic illustrating the functional interplay between BLM, SMARCAL1, and PRIMPOL in ALT-positive cancer cells and its therapeutic implications. In SMARCAL1-proficient ALT-positive cancer cells (left), replication stress at telomeric DNA—initiated and coordinated by BLM activity—generates ssDNA intermediates that are efficiently processed through the concerted action of SMARCAL1, FANCM, and the BLM helicase complex. This tightly regulated pathway sustains productive ALT activity, preserves telomere integrity, and supports cancer cell proliferation. In contrast, in SMARCAL1-depleted ALT-positive cancer cells (right), BLM-dependent telomeric stress persists but cannot be properly resolved, leading to uncontrolled ssDNA accumulation exacerbated by aberrant PRIMPOL-mediated repriming. This deregulated processing renders ALT non-productive, amplifies telomeric stress, and triggers robust DNA damage response activation. The resulting cellular outcomes include senescence or reduced cancer cell growth, thereby compromising tumor homeostasis. Senescence induced by SMARCAL1 depletion can be exploited through the use of senolytic agents to enhance tumor clearance.

## Notes

### Competing Interest Statement

The authors have declared no competing interest.

